# Tke5 is a novel *Pseudomonas putida* toxin that depolarises membranes killing plant pathogens

**DOI:** 10.1101/2025.01.11.632536

**Authors:** Carmen Velázquez, Alejandro Arce-Rodríguez, Jon Altuna-Alvarez, Jessica Rojas-Palomino, Andony Flores-Ceron, Citlaly Cando-Narvaez, Adrián Ruiz, Javier De la Peña Noya, Amaia González-Magaña, María Queralt-Martín, Antonio Alcaraz, David Albesa-Jové, Patricia Bernal

## Abstract

The soil bacterium *Pseudomonas putida* injects toxic proteins into neighbouring competitors, including resilient phytopathogens, using the Type VI secretion system (T6SS). The secretion of toxins endows *P. putida* with a significant fitness advantage, allowing this biocontrol agent to thrive in plant-related polymicrobial environments and prevent phytopathogen infections. Despite its agricultural significance, the toxin repertoire of *P. putida*, particularly those secreted via the K2- and K3-T6SSs, remains poorly understood. We present the first comprehensive molecular study of Tke5, a potent toxin encoded within the K3-T6SS, which represents the first comprehensive functional analysis of the BTH_I2691 protein family. Our biophysical data demonstrate that Tke5 is a pore-forming toxin that disrupts bacterial membranes through selective ion transport, inducing membrane depolarisation and cell death. Unlike conventional detergent-like pore-forming toxins, Tke5 preserves overall membrane integrity, avoiding large, non-specific disruptions. This unique mechanism offers a powerful approach to targeting resilient phytopathogens.

This study reveals a previously undescribed mode of action within a widespread yet understudied toxin family. Our findings highlight the potential of *P. putida* as a biocontrol agent, offering alternatives to chemical pesticides by exploiting novel toxin mechanisms. Understanding these bacterial toxins is crucial for developing effective strategies to combat plant pathogens.

## INTRODUCTION

Toxins are biochemical substances that can cause disease, death or cellular damage when introduced into or produced within an organism. They interfere with vital cellular processes such as inhibiting enzymatic activities or disrupting metabolism, protein biosynthesis or membrane functions. Among these diverse toxic substances, pore-forming toxins (PFTs) represent a particularly sophisticated class of molecular weapons that have evolved across multiple domains of life, being produced by various organisms, including bacteria, archaea, protozoans, vertebrates, and plants [1,2]. PFTs create pores in target cell membranes, leading to the influx or efflux of ions, small molecules, or proteins through the host membrane, which triggers various secondary responses, including cell death or more subtle manipulation of cell function [3]. PFTs are broadly classified into two categories based on the secondary structures that form the pore: α-PFTs use α-helices to form the transmembrane pore, such as colicins, cytolysin A (ClyA), and actinoporins; and β-PFTs utilise β-sheets/β-barrels to form the pore, which includes *Staphylococcus aureus* α-hemolysin, anthrax toxin or streptolysin O [1,3]. PFTs are soluble proteins that must undergo a transition from the soluble state into a membrane-spanning structure. They bind to the membrane of target cells via specific receptors or non-specific associations with the lipid bilayer. Upon membrane binding, PFT monomers oligomerise, inserting a specific pore-forming motif into the membrane to create a water-filled transmembrane pore. The structural complexity of these pores is evident in their stoichiometry and size, which can vary dramatically: from tetramers to structures involving 50 or more monomers. Consequently, pore dimensions range widely, from as small as 1-2 nm in small β-PFTs to more than 50 nm for large β-PFTs such as the cholesterol-dependent cytolysins (CDCs). The insertion process typically involves significant conformational changes in the protein structure, reflecting the dynamic nature of these molecular pore-forming mechanisms.

In bacterial contexts, toxins such as PFTs are often proteins secreted or injected into target cells as part of complex molecular mechanisms of defence or competition and are designed to target eukaryotic host cells or act against rival microorganisms, killing (bactericidal effect) or inhibiting the growth (bacteriostatic) of competing microorganisms. The type VI secretion system (T6SS) represents one of the best-characterised bacterial weapon systems, primarily deploying antimicrobial compounds against Gram-negative bacteria [4]. Its structural components include a cell wall-anchored membrane complex, a baseplate, a contractile sheath-tube assembly, and a spike. Effector proteins are loaded onto specific components, such as the inner Hcp tube and the VgrG-PAAR spike complex, and are ejected upon sheath contraction [5–11]. To ensure loading, T6SS effectors utilise specialised delivery mechanisms. Some effectors are encased in protective Rhs shells, while others rely on chaperones or adaptor proteins, like EagR and Tap. T6SS effector genes typically cluster near structural component genes (*vgrG, PAAR, hcp*) and are often accompanied by their chaperone and adaptor genes. Recent research has identified effector-specific motifs (MIX, FIX, RIX, PIX and WHIX) that are primarily N-terminal domains that facilitate the transport of the C-terminal toxic domain into target cells via the T6SS [12–17].

Recent studies have identified several T6SS-dependent PFTs, such as *P. aeruginosa* Tse5, which have a multi-ionic character, exhibiting a preference for cations while remaining permeable to anions [18,19]. Likewise, *P. aeruginosa* Tse4, known for its potent antibacterial activity, promotes cell depolarisation in competing bacteria by incorporating into cellular membranes and forming multi-ionic channels [20].

The production of protein toxins by bacteria for competitive advantage comes with a significant challenge: avoiding self-harm. To address this, toxin-producing bacteria have evolved a sophisticated self-protection mechanism. They produce specific immunity proteins that neutralise the effects of their own toxins, creating a crucial safeguard against self-intoxication. These immunity proteins are typically encoded immediately downstream of the toxin gene, forming a toxin-immunity pair. This genetic organisation serves two important functions: it ensures the simultaneous production of both the toxin and the cognate immunity protein, and it allows for the modular integration of the toxin-immunity pairs into diverse secretion systems [21]. While the exact mechanisms of immunity against PFTs are not fully understood, the immunity proteins for these toxins are usually small and hydrophobic, indicating that they likely localise to the cytoplasmic membrane. At this location, they may interfere with pore opening or disrupt the oligomerisation process of the pore-forming toxins [21].

The potent antibacterial activity of toxins has led to the exploration of their use in biological control. Many beneficial bacteria, such as some *Pseudomonas* species, act as effective biocontrol agents (BCAs) due to their ability to produce toxins that can inhibit or kill plant pathogens. Biocontrol is of significant importance in modern agriculture, offering a sustainable and environmentally friendly alternative to traditional chemical pesticides. Understanding the underlying mechanisms of toxins, including PFTs, could enhance the development of effective BCAs. In *P. putida*, a plant growth-promoting rhizobacteria (PGPR) well known for its biocontrol ability [22], the T6SS is instrumental in the elimination of severe plant pathogens [23]. *P. putida* KT2440 possesses three T6SSs (K1-, K2- and K3-T6SS) and their associated toxins [23], with predicted functions such as nucleases (Tke2, Tke4 and Tke10), pore-forming colicins (Tke6, Tke7, Tke8) or NAD^+^/NADP^+^ hydrolases, *i.e.* Tke1. However, no predicted functions exist for the putative Tke5, Tke9 and Tke3 effectors, and more importantly, none of the ten identified T6SS toxins of this strain have been characterised. Additionally, while the regulation of the K1-T6SS cluster has been thoroughly investigated [23,24], nothing is known about secretion through the K3-T6SS and its predicted associated effector Tke5. The primary reason for this lack of knowledge is that K3-T6SS is not functional under laboratory conditions [23,24], which limits the study of its components.

Advancing our understanding of novel toxins and their molecular mechanisms remains crucial for developing effective biocontrol strategies in sustainable agriculture. In this context, our work reveals the mechanism of action of Tke5, a putative T6SS effector of the biocontrol agent *P. putida,* providing the first biophysical characterisation of the BTH_I2691 protein family within the cl49522 superfamily. Tke5 forms ion-selective subnanometric pores in the cytoplasmic membrane of target cells, disrupting ion homeostasis that results in cell depolarisation without compromising membrane integrity. This discovery not only contributes to our fundamental understanding of bacterial competition, but the potent bactericidal activity of Tke5 against phytopathogens also opens new avenues for developing more effective and environmentally friendly crop protection strategies.

## METHODS

### Bacterial strains and growth conditions

Bacterial strains are listed in Table S1. Unless otherwise stated, chemicals and reagents, including antibiotics, were purchased from Sigma Aldrich. All strains were grown in Lysogeny broth (LB Lennox; 5 g L^−1^ NaCl) and agar (1.5% w/v) [25] for routine growth with shaking at 180 rpm, as appropriate. *E. coli* strains were incubated at 37°C, phytopathogens at 28°C and *P. putida* strains at 30°C. Vogel-Bonner Medium (VBM) (200 mg L^−1^ MgSO_4_.7H_2_O; 2 g L^−1^ anhydrous citric acid; 10 g L^−1^ K_2_HPO_4_; 3.5 g L^−1^ NaNH_4_HPO_4_.4H_2_O; pH 7.0) was used to select transconjugants after triparental conjugation, as *E. coli* donor and helper cells cannot grow using citrate as carbon source. Antibiotics were used at the following concentrations (µg mL^-1^): rifampicin (Rif), 20 for *P. putida*; kanamycin (Km), 50 for *P. putida*, *Agrobacterium tumefaciens*, *Pectobacterium carotovorum*, *Pseudomonas savastanoi*, *Pseudomonas syringae* and *Dickeya dadantii,* 25 for *E. coli* and *Ralstonia solanacearum,* and 20 for *Xanthomonas campestris*; ampicillin (Amp) 100 for *E. coli*, gentamycin (Gm), 25 for *P. putida* and 15 for *E. coli*; chloramphenicol (Cm), 10 for *E. coli;* nalidixic acid (Nal), 10 for *E. coli;* streptomycin (Sm), 50 for *E. coli*; tetracycline (Tc), 5 for *E. coli*.

### Construction of plasmids and bacterial strains

Plasmids and primers used in this study are listed in Tables S2 and S3, respectively. DNA manipulations were performed using standard methods [25]. Q5® High-Fidelity DNA polymerase (New England Biolabs) was used for PCR reactions according to the manufacturer’s instructions. Primers were synthesised by Macrogen Inc., and restriction enzymes were purchased from New England Biolabs. Plasmid isolation and purification of PCR products were performed with the Plus SV Miniprep DNA Purification System and the Gel and PCR Clean-Up System (Promega), respectively. All DNA constructs were sequenced and verified to be correct before use.

Recombinant plasmids were transferred to *E. coli* strains by transformation and to *Pseudomonas, Ralstonia, Agrobacterium, Dickeya, Pectobacterium, Erwinia* and *Xanthomonas* strains by electroporation [26] or conjugation [27], as appropriate.

The *tke5* (PP2612) gene was amplified from genomic DNA extracted from *P. putida* KT2440 using primers P1-P2 and P3-P2 (Table S3). Primer P3 is designed to fuse a *pelB* signal sequence at the 5’end. This sequence encodes the 22 amino acids, N-terminal leader peptide MKYLLPTAAAGLLLLAAQPAMA (UniProt Q04085) originally from the <u>Pe</u>ctate lyase <u>B</u> (PelB) of *Erwinia carotovora* EC [28]. PelB directs the fused protein, Tke5, to the Sec pathway [29]. The *tke5* and *pelB-tke5* genes were cloned into pJET1.2/blunt (Thermo Scientific™), sequenced and then subcloned into the broad host range, medium copy number vector pS238D•M (Table S2, [30]) from the SEVA collection at the NheI/BamHI sites, which excise the *msfGFP* to produce pS238D•*tke5* or pS238D•*pelB*-*tke5*. Primers P4-P5, that bind up- and downstream pS238D•M insertion site, were used to screen colonies harbouring the insert *pelB-tke5* and for subsequent sequencing.

The *tke5* gene was subcloned from pS238D•*tke5* to pCOLADuet^TM^-1 into *mcs-2*, including the TAA stop codon to avoid fusion with the C-terminal S-tag. Subsequently, a tag coding for 9x His residues and a Tobacco Etch Virus (-TEV) cleavage site was cloned upstream *tke5*, resulting in vector pCOLADuet-1•*9xhis-tke5*. These constructs were produced by GenScript service (Table S2).

The *tki5* (PP2611) gene, preceded by an artificial RBS region (TTTAA<u>AGGAGA</u>TATACAA), was synthesised and cloned into pSEVA424 vector between the KpnI and HindIII restriction sites using GenScript service (Table S2). The RBS and the *tki5* gene were subcloned into the broad host range, low copy number vector pSEVA624C at the KpnI/HindIII sites to produce pSEVA624C•*tki5* (Table S2). Vector pSEVA624C is a derivative of pSEVA621 from the SEVA collection [31,32] constructed in this study, where the *lacI*^q^-*P_trc_* cargo from pSEVA234C [33] was inserted in the *mcs* of pSEVA621 using PacI/HindIII. Primers P6-P7 were used to screen for the *lacI*^q^-*P_trc_* cargo insertion (Table S3) and primers P8-P9, which bind up- and downstream pSEVA624C *mcs*, to screen colonies harbouring the insert *tki5* and for subsequent sequencing. Vector pSEVA624C has been deposited into NCBI Genbank with accession number PQ628036 and into the SEVA repository https://seva-plasmids.com/ [31,32].

### Bioinformatic analysis

*Pseudomonas tke5* and *tki5* gene sequences and their respective protein product Tke5 and Tki5 were obtained from the Pseudomonas Genome database [34].

### Sequence homology search - Blast-P

BLASTP analyses were performed at the NCBI website using version 2.16.0 [35] and amino acid sequence searches using SMART [36] and Pfam [37] with the default settings.

### Conserved Domain Database - CDD

The CDD from NCBI was used to visualise the conserved domains of proteins Tke5 and Tki5. We accessed this database directly from the output of the BLASTP analyses. The CDD includes NCBI-curated domains, which use 3D-structure information to explicitly define domain boundaries and provide insights into sequence/structure/function relationships, as well as domain models imported from external source databases including Pfam, SMART, COG, PRK and TIGRFAMs [38].

### Protein Remote homology detection and 3D structure prediction - HHpred

Additionally, we have used the HHpred server to predict protein homology and 3D structure of Tke5 using the default parameters. This tool is part of the Max Planck Institute (MPI) Bioinformatics Toolkit.

### CDART

We used the Conserved Domain Architecture Retrieval Tool (CDART) from the NCBI website to visualise the different architectures present in the BTH_I2691 protein family. To do so, we introduce the accession number for this domain between brackets [NF041559] in the “Enter Query Protein Sequence” box under “Launch a new search” on the website of this tool. The output was downloaded in a comma-delimited table and is attached to this work as Dataset S1.

### Topology –TMHMM-2.0

The topology of Tke5 and Tki5 was predicted using the TMHMM-2.0 algorithm (DTU Health Tech; [39]) with the default settings (Supplementary Fig. 1).

### Phylogenetic tree

Protein sequences for phylogenetic analysis were retrieved from the NCBI website after the BLAST-P analysis. The multi-FASTA file was loaded in the Molecular Evolutionary Genetic Analysis (MEGA) tool version 11 [40] for alignment using the ClustalW algorithm. The resulting .mas file was exported as .mega to perform the evolutionary analysis by the Maximum Likelihood method with 500 bootstrap replications and JTT matrix-based model [41] using the same software (MEGA11), exact details as follows, as stated in the software after analysis:

“The evolutionary history was inferred by using the Maximum Likelihood method and JTT matrix-based model [41]. The tree with the highest log likelihood (−133685.63) is shown. Initial tree(s) for the heuristic search were obtained automatically by applying Neighbor-Join and BioNJ algorithms to a matrix of pairwise distances estimated using the JTT model, and then selecting the topology with superior log likelihood value. The tree is drawn to scale, with branch lengths measured in the number of substitutions per site. This analysis involved 71 amino acid sequences. There were a total of 1566 positions in the final dataset. Evolutionary analyses were conducted in MEGA11 [40]”.

The resulting file (.mtsx) was exported to a Newick file and loaded into iTOL software [42] for annotation and personalisation at the visualisation level.

### Structural-based homology search and protein interactions prediction - AlphaFold and Foldseek

The structure of Tke5 was predicted using AlphaFold2 and the PDB file was used as a query to search for homologs based on the structure using Foldseek Search Tool [43] with the default settings.

### Multiple Sequence Alignment (MSA) - Clustal Omega and Jalview

VasX from *V. cholerae* serotype O1 (strain ATCC 39315/ El Tor Iaba N16961) with the following identifiers: locus VC_A0020; Uniprot: Q9KNE5 VASX_VIBCH and Tke5 (PP2612) from *P. putida* KT2440 were aligned using the Multiple Sequence Alignment (MSA) tool of Clustal Omega (EMBL-EBI) [44] with the default settings and Jalview software was used to visualise the alignment (Supplementary Fig. 2). ChimeraX [45] was used for the visualisation of the AlphaFold predicted 3D structure of Tke5 and VasX.

### Growth inhibition and immunity-driven recovery assays

Cells of *P. putida* KT2440 were electroporated as previously described with a combination of the two compatible plasmids pS238D•M and pSEVA624C or their derivatives harbouring *tke5* or *pelB-tke5* and *tki5*, respectively, as described in Table S2. LB-agar supplemented with Km and Gm was used to select successfully transformed cells. The presence of the plasmid was corroborated by colony PCR using primers P4-P5 and P8-9 (Table S3) for pS238D•M and pSEVA624C derivatives, respectively.

To perform the growth curves, overnight cultures of each strain were grown in LB supplemented with the appropriate antibiotics. The next day, the cultures were adjusted to OD_600_ 0.05 in the same medium, and 1 mM of β-D-1-thiogalactopyranoside (IPTG) was added to induce *tki5*. Aliquots of 200 μL were placed in a microtiter plate and the Synergy/H1 microplate reader (Biotek©) was used to measure the OD_600_ every 15 min at 30 °C. After an initial incubation of 2 h, 0.5 mM of 3-methylbenzoate (3-*m*Bz) was added to each well and cultivated for another 14 h. Data was obtained for three biological replicates, each with technical duplicates.

### Survival growth in the presence of antimicrobial compounds

Overnight cultures of *E. coli* DH5α harbouring the vector pS238D•M [30] or the derivative encoding the PelB-Tke5 effector (pS238D•*pelB*-*tke5*, Table S2) were grown in LB medium supplemented with Km and diluted in 20 mL of the same medium at OD_600_ of 0.1. A 25 μL sample of each culture was taken at timepoint zero to quantify colony-forming units (CFU) as described below. When required, expression of *pelB*-*tke5* was induced with 0.5 mM 3-*m*Bz. A bactericidal (Gm_10_) or a bacteriostatic (Tc_5_) antibiotic was added to cultures of the control strain *E. coli* pS238D•M for reference to different substances affecting bacterial growth. The cultures were incubated at 30 °C/180 rpm for 6 h, 25 μL samples were taken every hour, and serial dilutions from 10^-1^ to 10^-6^ were prepared in a microtiter plate with phosphate buffered saline (PBS). A 20 μL aliquot of each dilution was placed onto 120 mm square LB-agar plates in technical duplicates. CFU were determined after 24 and 48h of incubation at 37 °C. The survival ratio was calculated based on the difference in CFU between each timepoint and timepoint zero, expressed as CFUt=x/CFUt=0. Cellular survival was represented as the Log_10_ of the survival ratio. Three biologically independent experiments were performed.

### Flow cytometry studies of E. coli DH5α cells expressing tke5, pelB-tke5, and tki5

Overnight cultures of *E. coli* DH5α alone or harbouring the vector pS238D (Calles et al., 2019), its derivative encoding the Tke5 (pS238D•*tke5*, Table S2) or PelB-Tke5 effector (pS238D•*pelB*-*tke5*, Table S2) and pSEVA624C•*tki5*, were grown in LB medium supplemented with Km and Gm when appropriate. Genes *tke5* and *pelB-tke5* were induced with 1 mM 3-*m*Bz when the cultures reached OD_600_ of 0.6 and incubated for 2 additional hours in the same conditions. Immunity gene *tki5* was induced with 1 mM IPTG for 1 hour before the 3-*m*Bz induction of the effector gene *tke5*. To avoid the leaking expression of the *P_trc_*, 0.2% glucose was added to the strains harbouring the pSEVA624C•*tki5* vector. Cultures were diluted in 10 mL of the same medium at OD_600_ of 0.05 and incubated for 2 hours once the 3-*m*Bz was added. Then, bacterial cultures were centrifuged at 6000 g for 15 min at 4 °C. To perform the flow cytometry studies, the bacterial pellets were resuspended in 1x PBS solution pH 7.4, to reach a final OD_600_ of 0.5.

To establish control populations, *E. coli* DH5α cells were subjected to heat shock at 85 °C for 5 minutes, followed by cooling at 4 °C for 2 minutes and stained with 4 μM of the fluorescent dye Sytox^TM^ Deep Red to identify the permeabilised population. For the depolarised population, *E. coli* DH5α cells were treated with the antibiotic polymyxin B sulfate (100 μg mL^−1^) for 30 min and stained with 10 μM of the fluorescent dye DiBAC_4_(3). Additionally, *E. coli* DH5α cells harbouring the pS238D vector, with and without 3-*m*Bz induction, were incubated with DiBAC_4_(3) and Sytox^TM^ Deep Red to serve as controls for the healthy population (Supplementary Fig. 3a-d). Cells were incubated with the fluorescent dyes for 45 min under constant shaking in dark conditions. Data was collected in the CytoFLEX cytometer with 488 and 638 nm lasers. Channels selected were FITC and APC for DiBAC_4_(3) (Exλ = 490 nm, Emλ = 516 nm) and Sytox^TM^ Deep Red (Exλ = 660 nm, Emλ = 682 nm), respectively.

*E. coli* DH5α strains harbouring either pS238D•*tke5* or pS238D•*pelB*-*tke5* or co-transformed with pS238D•*pelB-tke5* and pSEVA624C•*tki5* to assess the neutralisation effect of the immunity gene are shown in Supplementary Fig. 3e-h. Each measurement was performed in triplicate to determine the mean value, and each condition was tested in three-independent experiments (n=3). All data were subsequently analysed using CytExpert.

### Expression and purification of Tke5 to study pore-formation in lipid bilayers

*Escherichia coli* BL21(DE3) cells co-transformed with the pCOLADuet-1•*9xhis*-*tke5* and pSEVA624C•*tki5* plasmids were grown overnight at 37 °C with agitation in 200 mL of LB media supplemented with Km and Gm. For Tke5/Tki5 expression, bacterial cultures were diluted to an initial OD_600_ of 0.1 in 2 L of fresh LB medium supplemented with Km and Gm and induced when the OD_600_ reached 0.7 by adding IPTG to a final concentration of 1 mM. The temperature was lowered to 18 °C, and the bacterial cultures were left overnight in shaking conditions. Cells were harvested by centrifugation at 6000 × g for 20 min, and pellets were stored at −80 °C for subsequent use.

The pellet from 4 L of bacterial culture was resuspended in 60 mL of 50 mM Tris–HCl pH 8.0, 500 mM NaCl, 20 mM imidazole (solution A) along with 5 μL of benzonase endonuclease (Millipore, Sigma) and a tablet of protease inhibitor cocktail (cOmplete, EDTA-free, Roche). Cell lysis was performed using sonication with continuous cycles of 15 s ON and 59 s OFF at 60 % amplitude, followed by ultracentrifugation at 125,748 × g for 1 hour. The supernatant was filtered through a 0.2 μm membrane and then loaded via a peristaltic pump onto a 5-ml HisTrap HP column (GE Healthcare) equilibrated with solution A.

The column was connected to a fast protein liquid chromatography system (ÄKTA FPLC; GE Healthcare) and washed with solution A at 1 mL min^−1^ until the absorbance at 280 nm approached baseline. Tke5 was eluted using 100% solution B (50 mM Tris–HCl pH 8, 500 mM NaCl and 500 mM imidazole) at 3 mL min^−1^. The central peak fractions were combined, and dithiothreitol (DTT) was added to a final concentration of 2 mM. The protein solution was injected into a HiLoad Superdex 200 26/600 pg column, previously equilibrated with 20 mM Tris–HCl pH 8, 150 mM NaCl and 2 mM DTT (solution C). Tke5 eluted as a single monodispersed protein, appearing as a unique band on an SDS–PAGE gel (Supplementary Fig. 4). The identity of the Tke5 band was confirmed by mass spectrometry (Dataset S2). Peak fractions were pooled and concentrated using Amicon centrifugal filter units (30 kDa molecular mass cut-off, Millipore) to a final concentration of 2.5 mg mL^−1^ (*ca*. yield: 1.5 mg L^−1^) to perform lipid bilayer experiments.

### Study of Tke5 pore-forming activity in planar lipid bilayers

We used the solvent-free modified Montal-Mueller technique to enable the formation of planar lipid membranes in the orifice made over a Teflon film [46]. Lipid preparation involved chloroform evaporation of *E. coli* polar lipid extract (Avanti Polar Lipids; Alabaster, AL) under an argon stream, followed by dissolution in pentane at a concentration of 5 mg mL^−1^. The *E. coli* polar lipid extract headgroup composition consisted of 67% phosphatidylethanolamine (PE), 23.2% phosphatidylglycerol (PG), and 9.8% cardiolipin (CL), with acyl chains reflecting the intrinsic composition of *E. coli* [47].

A 20 µL aliquot of lipid solution was added to 1.8 ml of salt solution in both *cis* and *trans* compartments of the Teflon chamber, which was divided by a 15 μm thick Teflon film with a *ca*. 200 μm diameter orifice suitable for bilayer formation (Supplementary Fig. 5). Given the lipophobic nature of the Teflon film, lipid adherence was enhanced by pre-treating the orifice with a 3% hexadecane solution in pentane, which imparts a more lipophilic surface. After pentane evaporation, the salt solution levels in both compartments were raised above the orifice, facilitating planar bilayer formation through monolayer apposition. Capacitance measurements confirmed appropriate bilayer formation. Two distinct potassium chloride concentration gradients were employed for selectivity measurements: 250 mM KCl *cis* / 50 mM KCl *trans* (250/50 mM) and 50 mM KCl *cis* / 250 mM KCl *trans* (50/250 mM). For conductivity measurements, a symmetrical solution of 150 mM KCl *cis* / 150 mM KCl *trans* (150/150 mM) was used. All solutions were buffered with 5 mM HEPES at pH 7.4, with pH adjustment performed using a GLP22 pH meter (Crison). Tke5, dissolved in 20 mM Tris–HCl (pH 8), 150 mM NaCl, and 2 mM DTT, was added to the *cis* compartment to achieve a final concentration of 50 nM.

Ag/AgCl electrodes were custom-prepared within standard 250 μl pipette tips using 2 M KCl with 1.5% agarose bridges. A potential was considered positive when higher in the *cis* compartment (where the protein was always added), with the *trans* side set to ground. An Axopatch 200B amplifier (Molecular Devices, Sunnyvale, CA) in voltage-clamp mode facilitated current measurement and potential application. Collected data were filtered through an integrated 8-pole Bessel low-pass filter at 10 kHz and recorded with a sampling frequency of 50 kHz using a Digidata 1440A (Molecular Devices, Sunnyvale, CA). Data analysis was performed using pClamp 10 software (Molecular Devices, Sunnyvale, CA). For visualisation, current traces were digitally filtered with 500 Hz using a low-pass 8-pole Bessel filter.

The experimental setup, comprising a membrane chamber and head stage, was isolated from external noise sources by a double metal screen (Amuneal Manufacturing Corp., Philadelphia, PA). This configuration enabled measuring currents in the picoampere range with sub-millisecond time resolution. After potential application, the steady current was determined through Gaussian fitting of current value histograms. Currents were measured across a ± 0-100 mV range at 10 mV intervals, generating a single current-voltage (I-V) curve per experiment.

Ion selectivity measurements were conducted using concentration gradients between compartments of either 250/50 mM or 50/250 mM. Tke5-induced currents arise from the presence of at least one selective pore under a salt concentration gradient across compartments. The potential cation or anion preference of Tke5 currents was assessed by measuring the reversal potential (RP), defined as the applied voltage required to nullify the current.

A channel’s ion selectivity is determined by the RP value: a non-zero RP indicates selective ion transport. For a cation-selective channel, the RP would be negative under the 250/50 mM gradient and positive under the 50/250 mM gradient, with the opposite pattern applying to an anion-selective channel. Obtained RP values, derived from linear regression of measured current-voltage (I-V) curves, were corrected for liquid junction potential using Henderson’s equation to eliminate the salt bridge electrode contribution [48]. The corrected RP was subsequently introduced into the Goldman-Hodgkin-Katz (GHK) equation to calculate the channel permeability ratio (P^+^/P^-^) [49].

Channel conductance (G = I/V) was measured using a symmetric potassium chloride concentration of 150 mM/150 mM. Conductance increments (ΔG) were determined at ±100 mV, a voltage at which stable current increments or reductions were consistently observed. These ΔG values were calculated by averaging current increments (ΔI) and dividing by the applied voltage of ±100 mV. Normalised histograms of ΔG, generated with a 2.5 pS bin width, were fit to a single Gaussian distribution to determine the mean ΔG, which corresponded to the minimal channel conductance.

### Plant pathogens growth inhibition assays

Vector pS238D•M or the derivative encoding the PelB-Tke5 effector (pS238D•*pelB*-*tke5*) (Table S2) were transferred to cells of phytopathogens *P. syringae, R. solanacearum, A. tumefaciens, D. dadantii, P. carotovorum*, *E. amylovora*, *X. campestris* and *P. savastanoi* (Table S1) by conjugation as previously described. VBM agar supplemented with Km was used to select successfully transconjugants. The presence of the plasmids was corroborated by colony PCR using primers P4-P5 (Table S3). Overnight LB Lennox cultures of the abovementioned phytopathogens harbouring pS238D•M or pS238D•*pelB*-*tke5* were adjusted to an OD_600_ of 0.05 and 200 μL placed into a standard sterile 96 well-plate made of crystal-clear polystyrene with “U” form and 281 μL volume. The viability of the cells (OD_600nm_) was measured every 15 minutes for 22 hours at 28°C and 180 rpm using the Synergy/H1 microplate reader by Biotek©. Expression of *pelB-tke5* was induced after 2 hours with 0.5 mM of 3-*m*Bz except for *P. savastanoi,* where 0.1 mM was used. 3-*m*Bz was also added to the strains harbouring the *msfGFP* expressing vector pS238D•M to control the intrinsic toxicity of 3-*m*Bz in the absence of the effector. Data was obtained from three biological replicates, each with technical duplicates.

### Statistical analyses

Statistical analyses are based on Student’s *t*-tests in which two conditions are compared independently. P-values from raw data were calculated by two-tailed t-test and from ratio data to the control by one-sample t-test using GraphPad Prism 8 version 8.3.0.

For flow cytometry assays, the Brown-Forsythe and Welch ANOVA Test with Dunnett’s T3 multiple comparison test was performed to assess significant differences between the mean values of independent groups. Statistical significance was indicated as follows: not significant (ns), for p > 0.05, * for p ≤ 0.05, ** for p ≤ 0.01, *** for p ≤ 0.001.

## RESULTS

### Tke5 is a putative T6SS-dependent effector

The first study to characterise the T6SSs in *P. putida* identified ten putative T6SS effectors (Tke1 to Tke10) in the KT2440 strain [23]. The study did not determine the function of any of these effectors; although some were predicted to be nucleases (Tke2 and Tke4) or colicins (Tke6 and Tke7), the function of others, like Tke5, remained unknown [23]. Our current BLASTP analysis showed that Tke5 is a conserved T6SS component found in the Pseudomonadota phylum (formerly Proteobacteria), belonging to the BTH_I2691 family of the Conserved Domain Database (CDD), with accession number NF041559. A phylogenetic tree using the reference proteins of this NCBI family shows a wide distribution within the Beta- and Gammaproteobacteria classes (Fig. 1a). Most β-proteobacteria were from the Burkholderiales order, while γ-proteobacteria were more diverse, with many representatives from the Pseudomonadales order, but also Vibrionales and Enterobacterales (Fig. 1a). BTH_I2691 protein family is the only member of the recently created superfamily cl49522 (October 2023). In *P. putida* Tke5 effector, the BTH_I2691 domain spans amino acids 10 to 858 from a total of 996 amino acids (*E*-value 1.33e-117; Fig. 1b) according to the BLASTP result. Additionally, the CDD revealed a MIX_III domain of the MIX superfamily, also known as the VasX_N superfamily in the Pfam database, which is included within the BTH_I2691 domain. This domain was first identified as a Marker for Type VI effectors [12]. Curiously, it was named VasX_N by the Pfam database because it was present in the N-terminal region of the T6SS effector VasX from *V. cholerae*.

**Figure 1.**
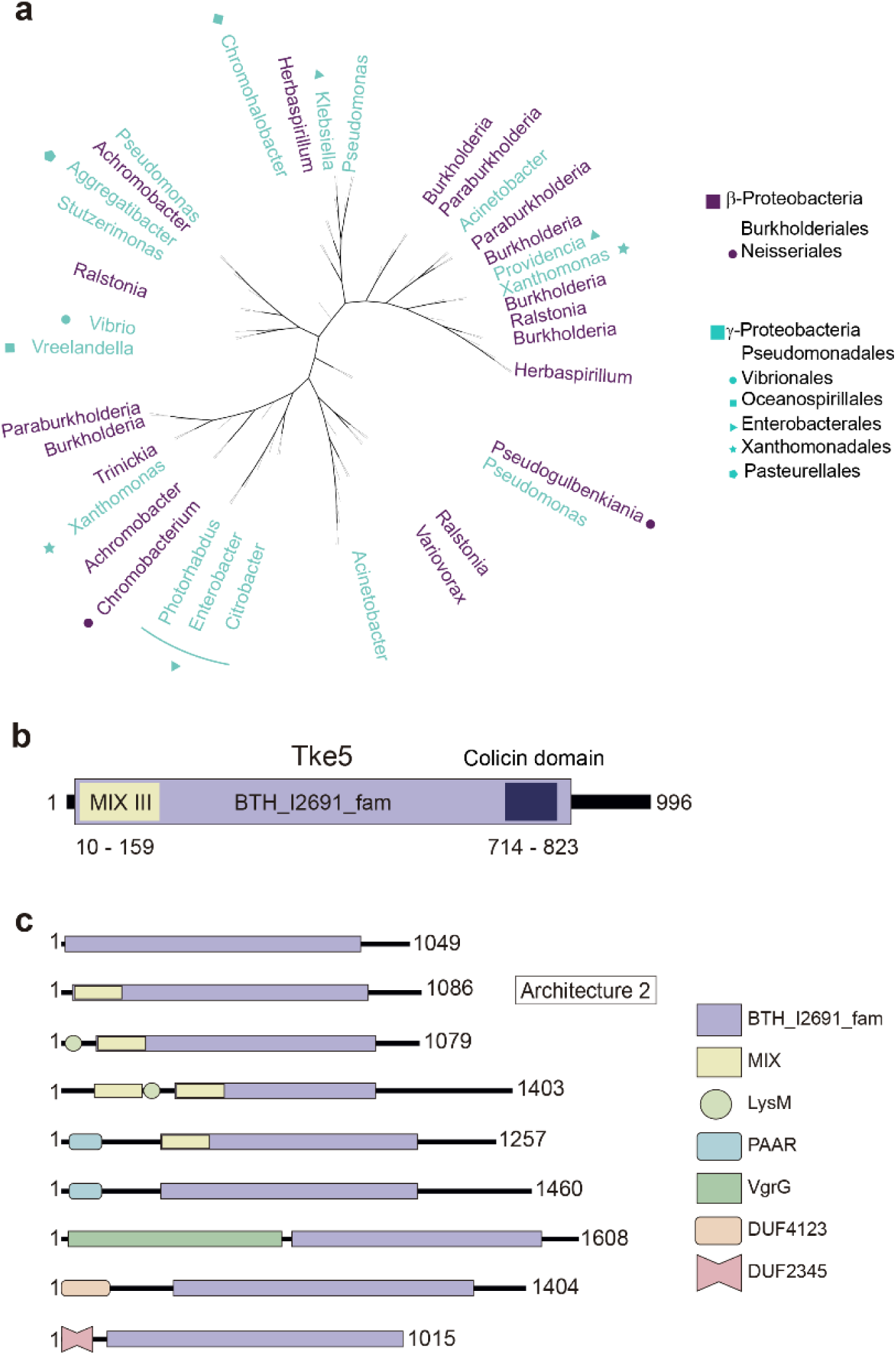
Tke5 *in silico* study. **a)** Phylogenetic distribution of BTH_I2691 proteins in the Pseudomonadota phylum. A maximum likelihood tree with 500 bootstrap replicates was built with Mega 11 for the reference proteins of the BTH_I2691 family. **b)** Prediction of conserved domains in Tke5 using BLAST-P analysis, showing the spanned amino acids along with the corresponding reference names and identification numbers. **c)** Conserved domain architecture of representative proteins within the BTH_I2691 family, indicating the amino acid spans and corresponding reference names.

VasX is the only member of the family previously identified. It was first described as a virulence factor secreted by *V. cholerae* T6SS necessary to kill the host model system *Dictyostelium discoideum* [50,51]. Subsequently, VasX was shown to be also an antimicrobial effector [51] which compromises the inner membrane of *Vibrio* species, and its activity is neutralised by the cognate immunity protein TsiV2 [50,51]. The effector presented a short domain (775-915 aa from 1085 aa) with homology to a pore-forming colicin also identified in Tke5 (Fig. 1b, dark blue), and preliminary results indicated that VasX compromised membrane integrity [52]. However, the exact mechanism of action of this effector is unknown.

The Conserved Domain Architecture Retrieval Tool (CDART) identified 61 different domain architectures within the BTH_I2691 protein family (Dataset S1), where the domain is found alone or in combination with many other domains. BTH_I2691 is mostly identified in the central part of these proteins (ranging from 1000 to 1500 amino acids), and interestingly, on many occasions, their N-terminal domains are also T6SS related. Fig. 1c shows representative proteins identified with CDART containing the T6SS-related domains *i.e.* MIX, PAAR, Tap (DUF4123), DUF2345 (VgrG related), LysM or even VgrG domains. Tke5 is most similar to Architecture number 2, which contains a BTH_I291 covering most of the protein and a single N-terminal MIX domain included in it (Fig. 1b).

To infer the function of this T6SS-related domain, we predicted the Tke5 topology that resulted in a 110-kDa membrane protein with five transmembrane domains (See Methods for details, Supplementary Fig. 1a). Moreover, we predicted the putative structure of Tke5 using AlphaFold 2 to perform a structural homology analysis with Foldseek [43]. This analysis retrieved that the Tke5 predicted structure was similar to the structure of *V. cholerae* toxin VasX from the AFDB Swissprot with an E-value of 9.82e-22, and the highest probability value (1) (Supplementary Fig. 1b), although the sequence identity was only 21% (Supplementary Fig. 2).

The genomic location of *tke5* within the K3-T6SS cluster (Fig. 2a), its structural homology to VasX (Supplementary Fig. 1b), and the presence of an N-terminal MIX domain associated with T6SS-dependent secretion and a C-terminal colicin domain (Fig. 1b) suggest that Tke5 may function as a T6SS-dependent toxin targeting the bacterial membrane.

**Figure 2.**
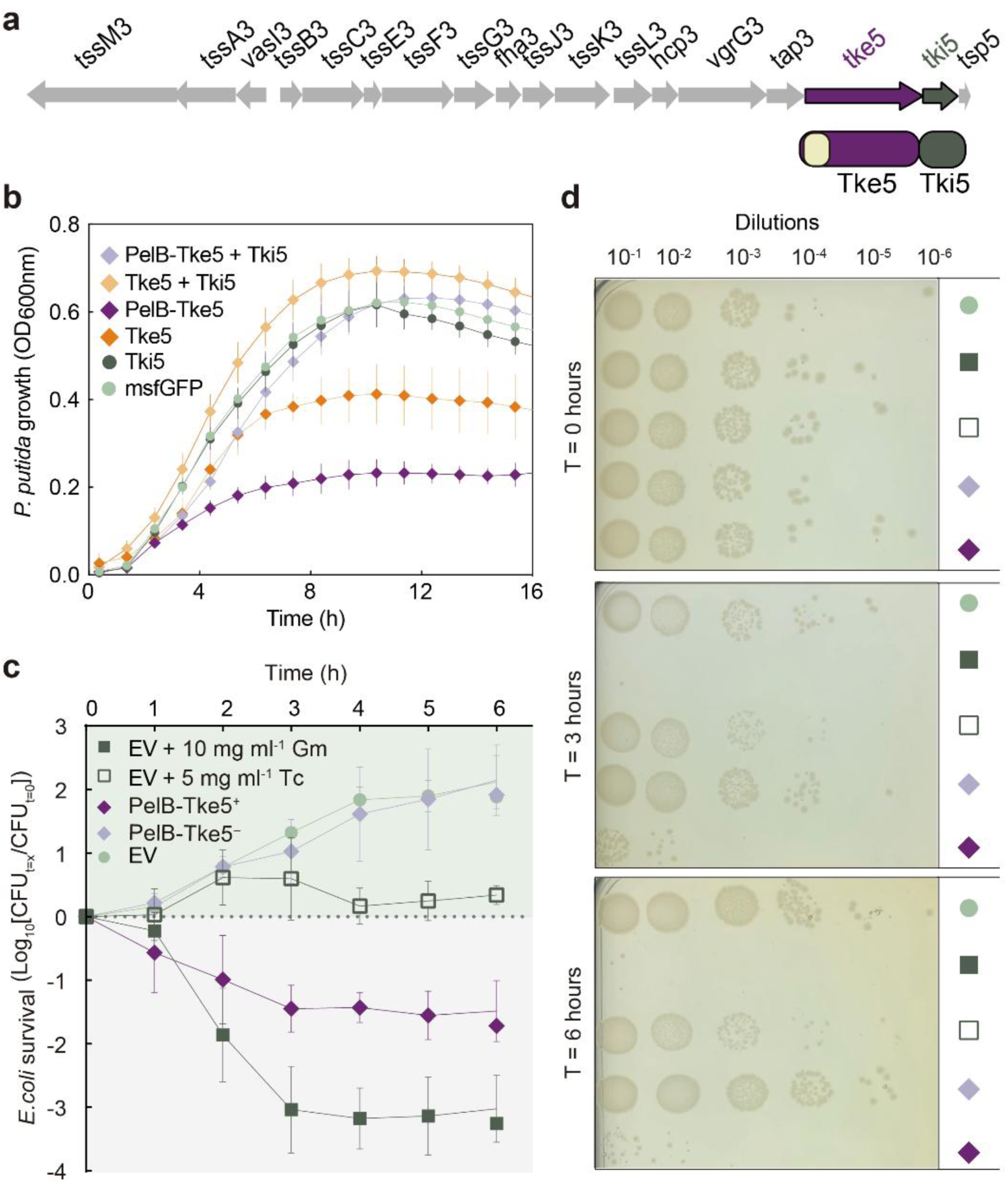
Tke5 interaction with its cognate immunity pair Tki5 can protect *P. putida* KT2440 from self-toxicity. **a)** Schematic representation of the genomic organisation of the genes of the K3-T6SS cluster from *P. putida*. Genes encoding core and accessory components are shown in grey, the *tke5* effector gene is depicted in purple with the MIX domain in yellow and the *tki5* immunity gene is coloured in green. **b)** *P. putida* growth curves under Tke5-Tki5 EI pair production. Overnight cultures were adjusted to OD_600_ 0.1, and the immunity protein was immediately induced with 1 mM IPTG. After 2 h of initial incubation at 30 °C, effector expression (*tke5* or *pelB-tke5*) was induced by the addition of 0.5 mM 3-*m*Bz (indicated with an arrow). OD_600_ was measured for a total of 16 hours. **c)** Bactericidal activity of Tke5. Survival rate curve of *E. coli* DH5α upon induction of *tke5* expression. CFU survival was quantified over 6 h after induction with 0.5 mM 3-*m*Bz. Gentamicin and tetracycline were included for comparison. Bacteria harbouring the *msfGFP*-bearing vector and the recombinant plasmid without induction were used as controls. 20 μL aliquots of each sample were taken hourly to determine the CFU ml^-1^ and calculate the Log_10_ survival rate (CFU_t=x_/CFU_t=0_). **d)** A subset (0, 3 and 6 hours) of each sample of *E. coli* DH5α cells from the different conditions tested in a) were plated in LB-agar to visualise the effect of Tke5 and the different controls on bacterial survival. The experiment was performed in triplicate.

### Tke5 is a potent antimicrobial toxin that can be neutralised by Tki5

To prove that Tke5 is an antimicrobial toxin, we tested the capacity of the putative Tke5 effector to impair bacterial growth when heterologously expressed from a plasmid. With this aim, the *tke5* gene was cloned into pS238D•M to produce pS238D•*tke5*. Additionally, a *pelB* sequence was fused to the *tke5* gene and cloned into pS238D•M to produce pS238D•*pelB*-*tke5*. The signal peptide PelB [53] was used to direct the effector to the Sec pathway to help a possible membrane insertion of Tke5, since the topology analysis revealed transmembrane helices, indicating that this effector might act within the bacterial membranes. The expression vector pS238D•M, known as the digitaliser (ON/OFF), contains a complex regulatory circuit that allows having an OFF state of zero transcription in the absence of induction and thus is ideal for toxin expression [30]. The plasmids pS238D•*tke5* and pS238D•*pelB*-*tke5* were introduced into *P. putida* KT2440, where the K3-T6SS is silent.

The expression of the effector gene was induced after two hours of bacterial growth in LB Lennox by the addition of 3-*m*Bz. Upon induction of the putative effector, the growth of the tested strain was significantly impaired, especially when fused to PelB (dark orange and purple curves, Fig. 2b), compared to the same strains harbouring the *msfGFP* reporter gene (light green curve, Fig. 2b). The expression of *msfGFP* was not deleterious, as expected for previous results [30]. These data indicate that Tke5 is a potent antimicrobial toxin (Fig. 2b).

Importantly, bacteria with active T6SS require immunity proteins to prevent self-intoxication from their own T6SS antimicrobial effectors and to protect against friendly fire from sister cells. Immunity genes are frequently located downstream of their cognate toxin genes to secure co-expression and thus protection. Therefore, we asked if the gene downstream *tke5, i.e., tki5,* codes for the immunity protein Tki5 (Fig. 2a). To assess the protective functionality of Tki5, the *tki5* gene was cloned into pSEVA624C, a vector compatible with pS238D (Fig. 2b), and then, *P. putida* was transformed with either pS238D•M, pS238D•*tke5* or pS238D•*pelB-tke5*. Expression of the immunity gene (*tki5*) was induced with IPTG at point zero, while the expression of the *tke5* toxin gene was induced with 3-*m*Bz two hours later. The co-expression of the putative *tki5* immunity gene together with the *tke5* or *pelB-tke5* rescued the impaired growth (light orange and purple curves, Fig. 2b). Although Tki5 is predicted to insert into the cytoplasmic membrane (Supplementary Fig. 1c), the expression of the *tki5* immunity gene in *P. putida* KT2440 does not exhibit toxicity (dark green curve, Fig. 2b), which is expected since KT2440 is the natural host of this protein. These data show that Tki5 protects *P. putida* KT2440 against Tke5 intoxication, demonstrating that Tke5/Tki5 are an effector-immunity (EI) pair.

### Tke5 is a bactericidal antimicrobial toxin

Here, we have demonstrated that Tke5 is a powerful antimicrobial toxin. Depending on their effect on bacterial targets, antimicrobial compounds can be classified as bactericidal if they kill bacteria, or bacteriostatic if they inhibit bacterial growth without directly killing the bacteria (*i.e.*, induce growth arrest). To determine whether Tke5 acts as a bactericidal or bacteriostatic agent, we performed cellular survival assays under the expression of *tke5* using the laboratory host cell *E. coli* DH5α. Two antimicrobial substances with known mechanisms of action were used as controls: the bactericidal antibiotic gentamicin (Gm) and the bacteriostatic antibiotic tetracycline (Tc). Both target the 30S ribosomal subunit binding to the 16S rRNA, but their mechanisms differ significantly. Gm binds irreversibly, fooling the ribosome and leading to indiscriminate tRNAs acceptance, which results in defective proteins and ultimately cell death. In contrast, Tc binds reversibly and prevents the peptide chain elongation. We have represented the cellular survival as the Log_10_ of the survival rate, which is the ratio between the CFU at a given time and the CFU at time zero (Log_10_ survival rate [CFU_t=x_/CFU_t=0_]). Thus, the cellular survival of an organism growing in the absence of an antimicrobial compound will have a value greater than 0 (*E. coli* harbouring the empty plasmid in the absence of antimicrobial compound, light green curve, Fig. 2c). In the presence of a bacteriostatic agent, the CFU count remains constant over time, resulting in a survival rate of 1 and thus the Log_10_ representing cellular survival is 0 (*E. coli* harbouring the empty plasmid in the presence of Tc, dark green empty square curve, Fig. 2c). Conversely, a microorganism exposed to a bactericidal agent will present a cellular survival below 0 (*E. coli* harbouring the empty plasmid in the presence of Gm, dark green full square curve, Fig. 2c). Upon induction of *tke5*, *E. coli* survival at every tested point (hourly for a total of 6 hours) was consistently below 0 as a bactericidal agent (dark purple diamond curve, Fig. 2c), while the uninduced strain (light purple diamond curve, Fig. 2c) reached a cellular survival above 2, showing growth similar to the strain with the empty plasmid.

Serial dilutions of these cultures were plated on LB agar plates at points 0, 3 and 6 hours to check survival (Fig. 2d). Individual CFU from cultures grown in the absence of antimicrobial compounds were visualised in the dilutions 10-e4 and 10-e5. As expected, the presence of Tc slightly decreased growth but did not kill the population, while no CFU could be visualised in the cultures treated with Gm (Fig. 2d). Confirming the growth curve survival assay, the bactericidal effect of Tke5 was evident at 3 hours upon gene induction, while the non-induced culture grew at the same level as the control strain (Fig. 2d). All this data indicates that Tke5 is a potent bactericidal toxin.

### Tke5 induces cell depolarisation that can be neutralised by Tki5

Our data support that Tke5 is a potent bactericidal toxin (Fig. 2b-d) whose toxicity is neutralised by the Tki5 immunity protein (Fig. 2b). Additionally, our bioinformatic analysis suggested that Tke5 could be targeting the bacterial membrane (Supplementary Fig. 1a).

To evaluate this possibility, we measured the effect of Tke5 on cell permeability and membrane potential of intoxicated *E. coli* DH5α cells using flow cytometry. To this end, *E. coli* cells were transformed with the plasmid coding for Tke5 (pS238D•*tke5*) or the variant encoding Tke5 fused to the PelB leader sequence (pS238D•*pelB*-*tke5*). Cells were cultured in liquid media, and the expression of *tke5* or *pelB-tke5* was induced with 3-*m*Bz. Two hours after *tke5* induction, cells were stained with Sytox^TM^ Deep Red and DiBAC_4_(3) to measure cell permeability and cell depolarisation, respectively. Sytox^TM^ Deep Red is a nucleic acid stain that penetrates cells with compromised membranes, resulting in red fluorescence, thus allowing the detection of permeabilised cells by flow cytometry [54]. DiBAC_4_(3) is an anionic probe that binds to intracellular proteins or the membrane upon permeation into cells that have lost their resting membrane potential and become depolarised, resulting in green fluorescence [55].

To define the possible cell populations resulting from the abovementioned treatment, *i.e.* healthy, permeabilised and depolarised cells, we performed the following controls: i) cell permeabilisation was induced by heat shock (red fluorescence, top left square, Supplementary Fig. 3a), ii) cell depolarisation was induced by treatment with polymyxin B antibiotic (green fluorescence, bottom right square, Supplementary Fig. 3b), and iii) untreated cells harbouring the empty pS238D plasmid uninduced and induced with 3-*m*Bz, represented the healthy population (mostly non-fluorescence, bottom left square, although *ca.* 7% showed green fluorescence, bottom right square, Supplementary Fig. 3c and 3d, respectively).

Measurements of cells expressing *tke5* and treated with Sytox^TM^ Deep Red and DiBAC4(3) revealed that the production of Tke5 and PelB-Tke5 results in an approximate reduction of 5% and 16% in the healthy cell population, respectively, compared to the negative control (Fig. 3a, Supplementary Fig. 3e and f). These reductions in the healthy population corresponded with an increase in depolarised cells upon Tke5 and PelB-Tke5 induction (Fig. 3b, Supplementary Fig. 3e and f), while the percentage of permeabilised cells (Fig. 3c) or permeabilised and depolarised cells (Fig. 3d) remained unchanged and near-zero.

**Figure 3.**
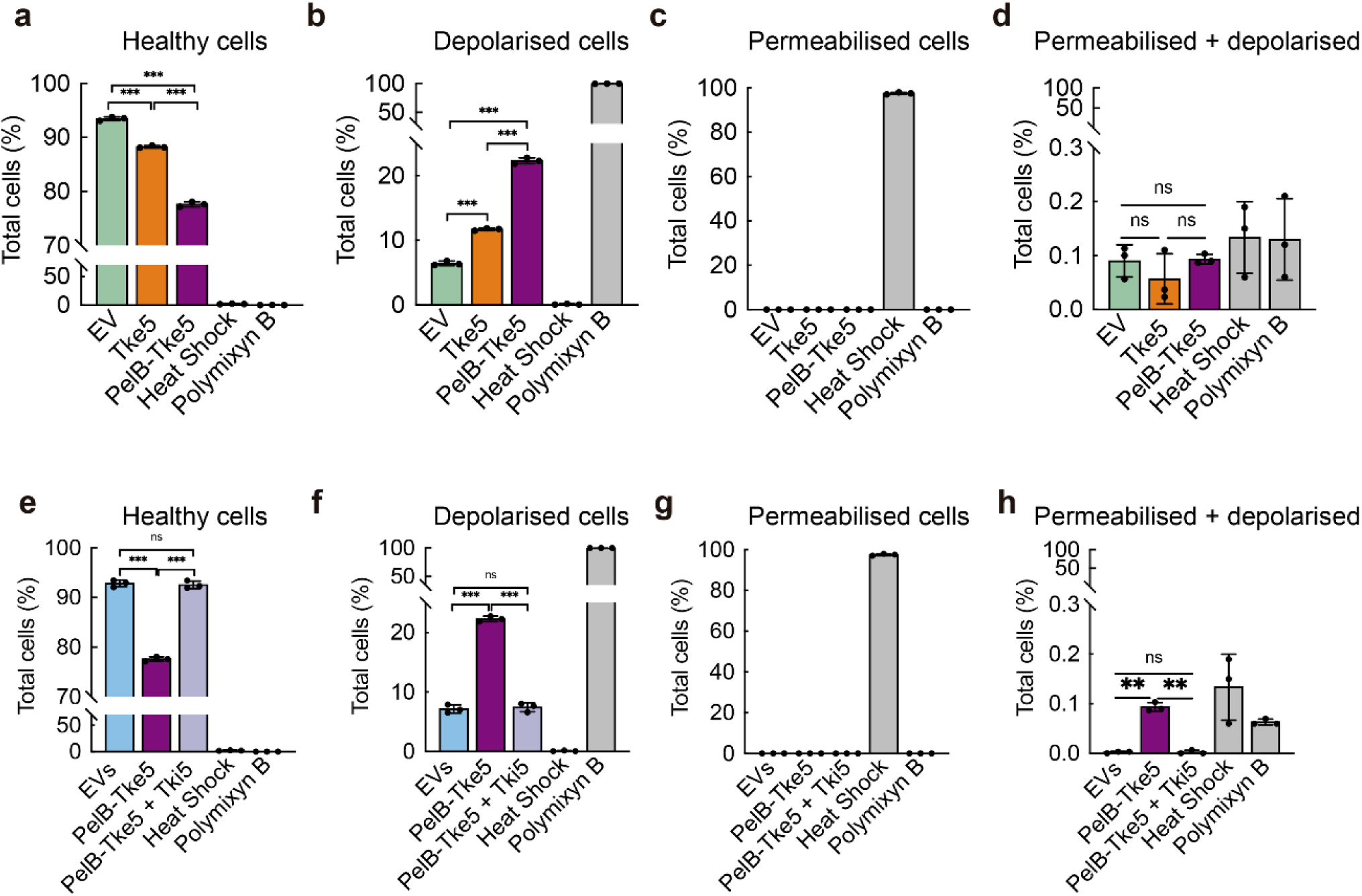
Cell depolarisation induced by Tke5 effector and toxicity protection from Tki5 immunity protein. Flow cytometry assays were carried out in *E. coli* DH5α cells transformed with the corresponding vectors and stained with both DiBAC4(3) (depolarised population) and Sytox^TM^ Deep Red (permeabilised population). The upper row represents the percentage of healthy (**a**), depolarised (**b**) permeabilised (**c**) and permeabilised + depolarised (**d**) cell populations upon production of Tke5 effector with and without being fused to PelB. Cells with the empty vector, subjected to heat shock and treated with polymyxin B are included as control samples. The bottom row represents the percentage of healthy (**e**), depolarised (**f**), permeabilised (**g**) and permeabilised + depolarised (**h**) cell populations upon expression of PelB-Tke5 effector with and without the co-expression with the immunity protein Tki5. Cells without induction, subjected to heat shock and treated with polymyxin B are included as control samples.

These results indicate that Tke5 and PelB-Tke5 alter the membrane potential of intoxicated cells without affecting the integrity of the cytoplasmic membrane. Consistent with our previous data (Fig. 2b), the impact of Tke5 is increased in the presence of the signal peptide PelB (Fig. 3b). As expected, based on results where Tke5 toxicity is neutralised by Tki5 (Fig. 2b), co-expression of PelB-Tke5 with the immunity protein Tki5 prevents cell depolarisation (see Methods for details; Fig. 3e-h).

Given that our results suggest that Tke5 targets the cytoplasmic membrane of intoxicated cells, causing membrane depolarisation, we hypothesised that Tke5 may be a pore-forming protein that allows ion transport across the membrane, destabilising the resting membrane potential. This ion transport could represent the molecular mechanism employed by Tke5 to depolarise intoxicated cells.

### Tke5 forms ion-selective pores in model membranes

To examine the ion channel activity of Tke5, we expressed and purified the protein from the cellular soluble fraction of *E. coli* BL21(DE3) cells co-transformed with pCOLADuet-1•*9xhis-tke5* and pSEVA624C•*tki5* plasmids (see Methods and Supplementary Fig. 4 for details) and evaluated its properties in a model membrane. Importantly, Tke5 is soluble in aqueous solution, exhibiting a dual state, either soluble or membrane-inserted, similar to pore-forming colicins. Using a modified solvent-free Montal-Mueller technique [46], we formed a synthetic lipid bilayer covering a small aperture separating two chambers (*cis* and *trans*) connected to an electric circuit (Supplementary Fig. 5). These chambers, filled with buffers of defined salt concentrations, provide a well-controlled artificial environment using only minimal amounts of material. In contrast to liposomes, one has direct access to both sides of the bilayer and, therefore, it is possible to work in asymmetric conditions in terms of electrolyte and membrane composition. Following protein addition, we can measure ionic currents passing through Tke5-induced membrane pores. This experimental setup allows us to assess the pore conductance and ionic selectivity (cation *versus* anion preference).

To start, we assembled the synthetic lipid bilayer using an *E. coli* polar lipid extract in 150 mM KCl and confirmed membrane impermeability by ensuring the absence of leakage current. Then we added Tke5 to the *cis* chamber at a final concentration of 50 nM and waited for protein insertion until we observed stable currents indicative of ion flow through Tke5-formed pores. Fig. 4a shows a representative current trace (current intensities denoted as I) as a function of applied potential (V). The I/V relationship was linear at low voltages, indicating ohmic behaviour (Fig. 4b, circles), while higher voltages produced small stepwise current jumps (insets in Fig. 4a). Currents in between jumps remained linear (Fig. 4b, squares and diamonds), indicating that the ohmic behaviour was maintained at higher voltages and after the appearance of current jumps. Of note, the frequency of these random transitions between conducting levels increased when the voltage surpassed *ca.* 70 mV, indicating either a change in the number of Tke5 pores inserted into the lipid bilayer or, alternatively, dynamic structural changes in the pore size of already existing Tke5 channels.

**Figure 4.**
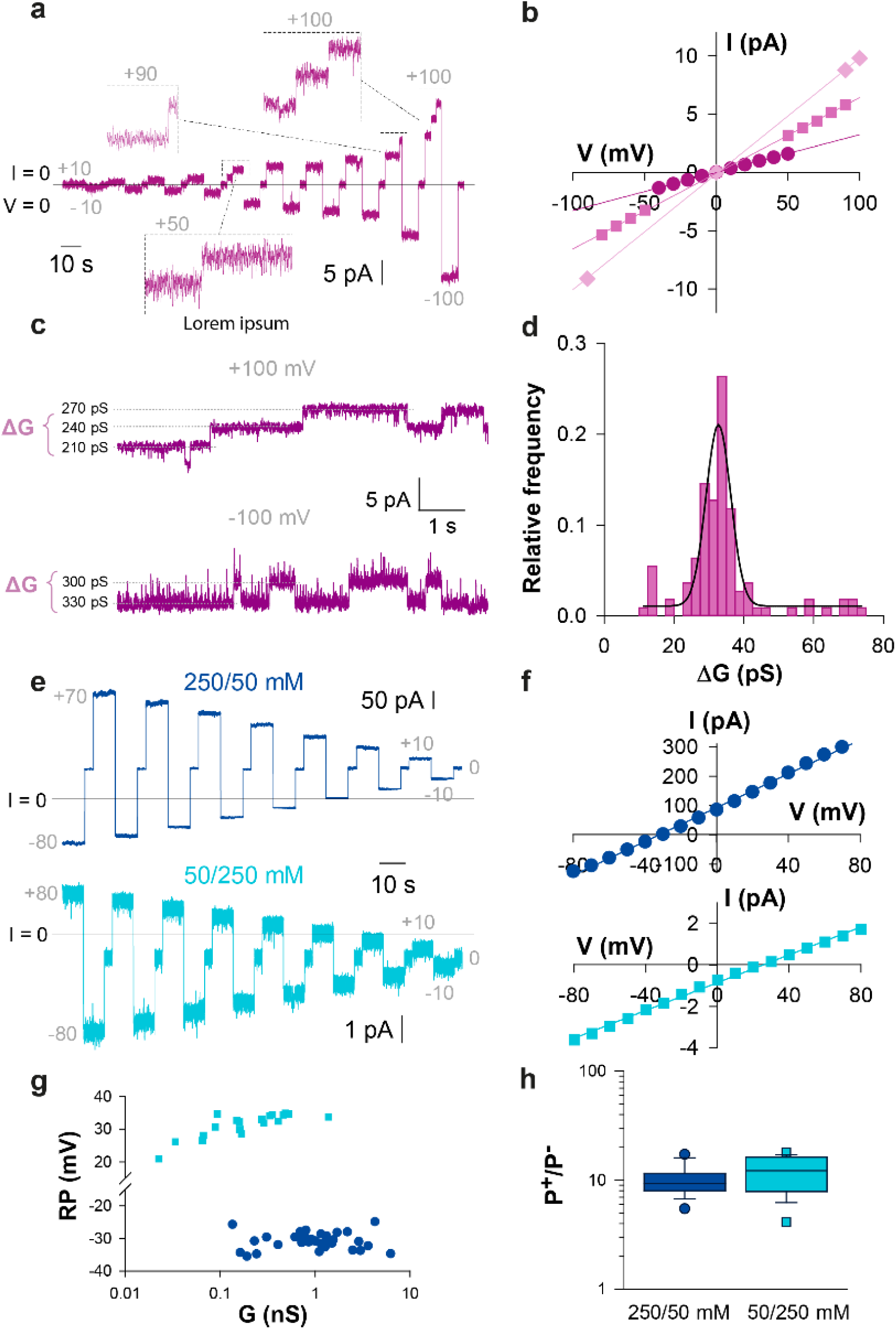
Tke5 forms stable, cation-selective pores in model membranes. **a)** Representative current trace induced by Tke5 under symmetric 150/150 mM KCl condition. The horizontal black line marks zero applied voltage and zero current. Voltage was applied starting at ±10 mV and increasing it in 10 mV steps up to ±100 mV. Some voltages are indicated in grey. Dashed-lined rectangles highlight current increments at +50, +90 and +100 mV. This trace is representative of 24 independent experiments. **b)** Current-voltage (I/V) plot corresponding to the trace in (a). Solid lines correspond to linear fittings of the experimental data with R² = 0.994 (circles) and R² = 0.9999 (squares and diamonds). **c)** Representative current traces induced by Tke5 under symmetric 150/150 mM KCl and recorded while maintaining a ±100 mV voltage for long time. Dotted lines indicate the conductance of each level, which are used to calculate the conductance increments ΔG = ΔI / V, as indicated. This trace is representative of 18 independent experiments. **d**) Normalised histogram (bin of 2.5) showing conductance increments ΔG obtained at ± 100 mV in 150/150 mM KCl as those shown in panel (c). Values are fitted to a Gaussian, with the mean ΔG representing the most probable channel conductance jumps. Histogram built from 110 different current jumps obtained in 18 independent experiments. **e)** Representative stable current traces induced by Tke5 in asymmetric 250/50 mM and 50/250 mM KCl conditions, shown in dark and light blue, respectively. Voltage was applied starting at −80 mV (250/50 mM) or at +80 mV (50/250 mM) and decreased in 10 mV steps down to ±10 mV. Some voltages are indicated in grey. The horizontal black line indicates zero current. These traces are representative of 32 (250/50 mM) and 19 (50/250 mM) independent experiments. **f)** I/V plots for traces in (e). **g)** Plot of RP (mV) versus conductance (nS) for independent measurements in 250/50 mM (n=33) and 50/250 mM (n=19) KCl conditions. A negative RP in 250/50 mM KCl gradient condition and a positive RP value in the 50/250 mM KCl gradient condition indicate Tke5 cation selectivity. **h)** Box plot showing cation vs. anion permeability ratios (P^+^/P^-^) derived from RP values in (**g**). Boxes display the 25^th^ and 75^th^ percentiles, with a line at the median and error bars defining the 10^th^ and 90^th^ percentiles.

To distinguish between these two possibilities, we first estimated the size of the Tke5 pore lumen by applying ±100 mV voltage pulses for long durations and measuring Tke5-induced current jumps (ΔI) (Fig. 4c). We then analysed their respective conductances (ΔG=ΔI/V), which reflect their current-carrying capacity. Histogram in Fig. 4d displays the values obtained and shows a clear peak centred around ΔG = 33 ± 4 pS, which aligns with the known conductance in the same salt and lipid conditions of Tse4, another effector secreted by T6SS and recently demonstrated to form narrow ion-selective membrane pores [20]. Tse4 structure is not yet known, but a radius of ∼0.5 nm has been estimated for Tse4-induced pores based on electrophysiology data and previous studies showing that small solutes between 300-700 Da (corresponding to 0.45 – 0.65 nm equivalent hydrodynamic radius) could not access the cytoplasm of Tse4-intoxicated cells [20,56]. Thus, given the comparable conductance of Tke5 and Tse4 (33 pS vs. 20 pS), we can infer a radius of less than 1 nm also for Tke5. Such a subnanometric pore diameter is typical of highly specialised narrow channels allowing only the transport of partially dehydrated ions while excluding larger hydrated ions, other solutes and metabolites [57].

The broad conductance range (10-80 pS) observed in Fig. 4d may reflect some variability in the size of Tke5 pores formed among experiments or also the presence of multiple simultaneous insertions operating as clusters. To address this issue, we assessed the ionic selectivity of Tke5-induced pores by measuring the reversal potential (RP) and compared it with the conductance data. The RP is the potential needed to null the current flow through the channel under asymmetric salt conditions [58,59]. Since the physical dimensions of the channel lumen define the intensity of the interaction between the pore charges and the permeating ions, the ionic selectivity of the channel increases as the channel becomes narrower, and vice versa. Accordingly, if the RP necessary to counteract the flow of ions does not change significantly with G, this would mean that a number of quasi-identical channels are inserted simultaneously in the membrane. In contrast, a decrease in RP with increasing G would indicate a size-dependent population of channels [59,60].

To study the RP of Tke5 channels, we measured induced currents at two concentration gradients: first with 250 mM KCl in the *cis* chamber and 50 mM KCl in the *trans* chamber (250/50 mM KCl), and then with an inverted gradient of 50/250 mM KCl, (dark blue/light blue data represented in Fig. 4e-h). The gradient mimics the ion concentration differences across bacterial membranes, where total ion concentrations are higher in the periplasm than in the cytosol as a result of charged macromolecules being unable to cross the inner membrane. Representative current traces with the 250/50 mM and 50/250 mM KCl gradients are shown in Fig. 4e. For each experimental trace, we plotted the I/V curve (Fig. 4f) to confirm the ohmic behaviour and calculated the conductance associated with each RP measurement (Fig. 4g). In the 250/50 mM KCl gradient, the Tke5 currents produced values of RP ∼ 30 mV which is similar to the absolute RP obtained in the 50/250 mM KCl gradient ∼ −30. The fact that RP does not change significantly in a variety of Tke5 channels displaying different G values (several orders of magnitude) suggests the presence of clusters of channels with similar dimensions adding to the total ion flow, rather than the assembly of larger pores, which would have altered considerably the RP values in Fig. 4g.

The RP also reveals channel selectivity, *i.e.* its relative preference for cations *versus* anions. When a channel favours cations, more positive ions move through the channel from the side with a higher concentration to the side with a lower concentration. Therefore, when the salt concentration is higher on the *cis* side and lower on the *trans* side (250/50 mM KCl gradient), the RP will be negative to counteract the movement of the cations from *cis* to *trans* chambers (note that in our set up the *trans* compartment is grounded, see Methods and Supplementary Fig. 5). Conversely, if the gradient is inverted, the RP will be positive. This is exactly what we observed in our setup, where the average RP for the 250/50 mM KCl gradient is negative (−31±2 mV), while the average RP for the 50/250 mM KCl gradient is positive (+33±3 mV) (Fig. 4g), indicating that Tke5 channels preferentially allow positively charged ions (such as K^+^) to flow through.

Lastly, to obtain a quantitative estimation of the ratio between the flow of cations and anions, *i.e.* the ion permeability ratio (P_K_^+^/P_Cl_^-^), we used the Goldman-Hodgkin-Katz flux equation (GHK, [61]), which yielded values of P_K_ ^+^/P ^-^ = 10 ± 3 and 16 ±9 in the 250/50 and 50/250 mM KCl gradient, respectively (Fig. 4h), (note that GHK equation shows an exponential relationship between P_K_ ^+^/P ^-^ and RP, so that small changes in RP yield noticeable differences in P_K_ ^+^/P ^-^). Weakly selective channels tend to have a P_K_ ^+^/P ^-^ <5, while ideal selective channels can reach values of 100-1000 [57,59]. Therefore, these results indicate that Tke5 forms subnanometer ionic channels with a relatively high preference for K^+^ over Cl^-^ ions.

### The Tke5 toxin kills plant pathogens

*P. putida* is a biocontrol agent that uses the T6SS to kill plant pathogens *i.e.* phytopathogens[23]. Thus, we tested the capacity of the putative Tke5 effector to kill phytopathogens of importance for agriculture. We selected a list of pathogens that included at least one strain of most of the Top 10 plant pathogenic bacteria species based on scientific and economic importance [62] *i.e.,* (1) *Pseudomonas syringae* pathovars; (2) *Ralstonia solanacearum*; (3) *Agrobacterium tumefaciens*; (4) *Xanthomonas oryzae*; (5) *Xanthomonas campestris*; (6) *Xanthomonas axonopodis*; (7) *Erwinia amylovora*; (8) *Xylella fastidiosa*; (9) *Dickeya dadantii*; (10) *Pectobacterium carotovorum*, and (11) *Pseudomonas savastanoi*.

Among the crops affected by these phytopathogens are brassica crops including broccoli, cauliflower, cabbage, turnip, and mustard (*X. campestris*); coffee, kiwi, tomato, rice, beans, cherry, plum, horse chestnut, loquat (*P. syringae*); carrot, potato, leafy greens, squash, onion, peppers (*P. carotovorum*); walnuts, grapevines, stone fruits, nut trees, sugar beets, horseradish, rhubarb (*A. tumefaciens*) and olive trees, oleander, and ash trees (*P. savastanoi*); the main causal agents are indicated in brackets immediately after the affected crops.

To test the efficiency of Tke5 in impairing the growth of plant pathogens, the broad-host range plasmid pS238D•*pelB*-*tke5* was transferred by conjugation to the strains of interest *i.e.*, *P. syringae* pv. syringae B728a, *R. solanacearum* NCPPB 1493, *A. tumefaciens* C58, *D. dadantii* 3937, *P. carotovorum* subsp. carotovorum SCRI 194, *E. amylovora* NCPPB 595, *X. campestris* pv vesicatoria NCPPB 195 and *P. savastanoi* pv. savastanoi NCPPB 3335. The expression of the effector gene was induced after two hours of bacterial growth in LB Lennox by the addition of 3-*m*Bz. Upon induction of the putative effector, the growth of the different tested plant pathogens was significantly impaired in all cases (dark purple curve, Fig. 5) compared to their controls, that is, the same strains harbouring the pS238D•M vector (green curves, Fig. 5) or without 3-*m*Bz induction (light purple curve, Fig. 5). The addition of 3-*m*Bz to the plant pathogen strains harbouring the pS238D•M vector, which expresses the *msfGFP* reporter gene instead of the *tke5* gene effector, demonstrated that: i) the concentration of 3-*m*Bz used in our assay is not toxic to these strains, and ii) the level of expression of this system is not inherently toxic, as the production of a non-toxic protein, msfGFP, did not affect the survival of these strains (dark green curve, Fig. 5).

**Figure 5.**
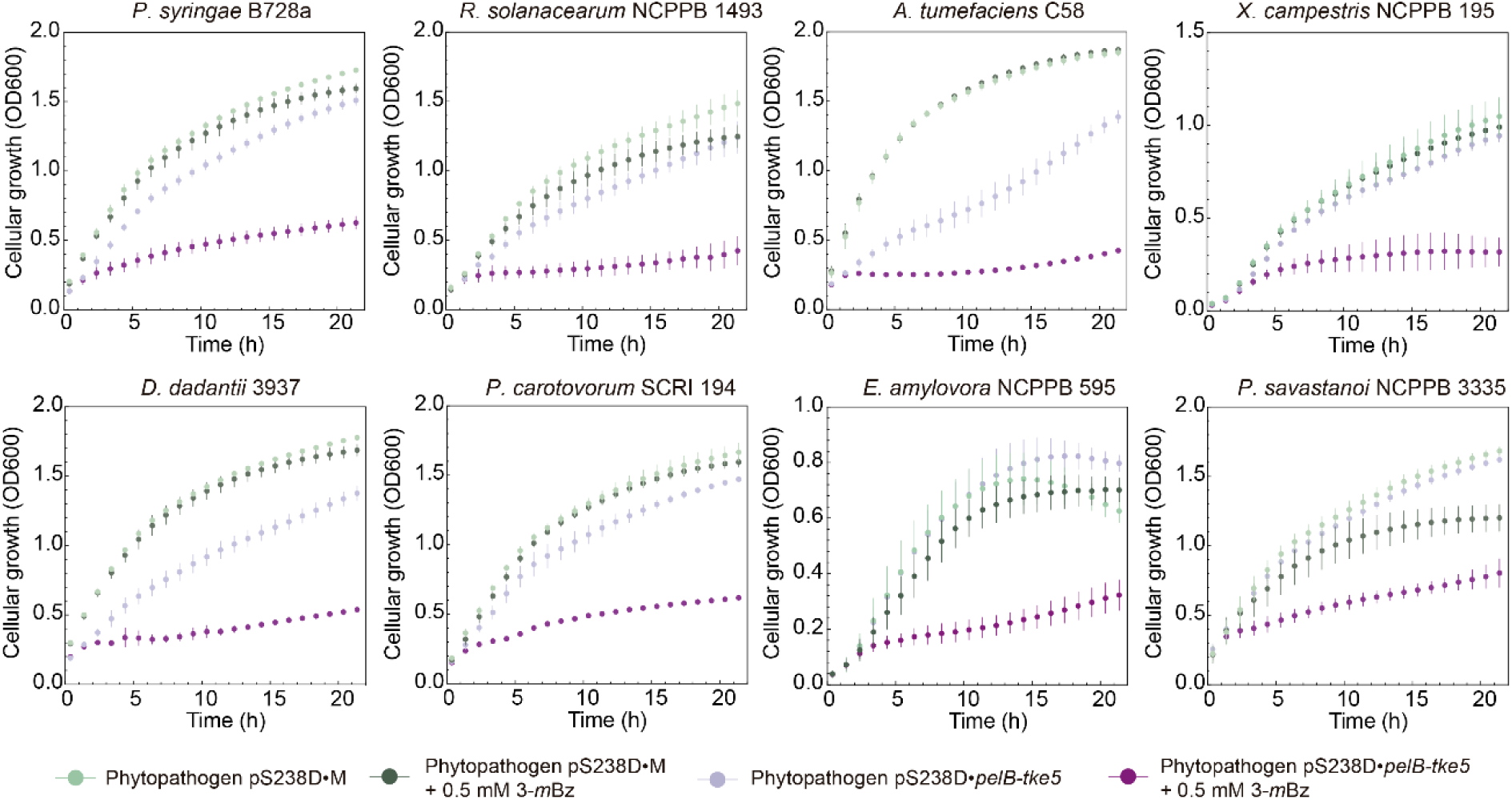
Toxicity of Tke5 for different recalcitrant phytopathogens. The growth of plant pathogens cells harbouring the pS238D•*pelB*-*tke5* containing the T6SS effector of *P. putida* Tke5 N-terminal fused to a PelB signal peptide was determined by measuring the OD at 600 nm every 15 minutes for 22 hours at 28°C using the Synergy/H1 microplate reader by Biotek©. After 2 hours, 3-*m*Bz was added to the LB medium to induce the expression of the *pelB-tke5* gene.

All these data demonstrate the potent antibacterial activity of the Tke5 T6SS toxin against a diverse range of economically significant plant pathogens. The broad-spectrum inhibitory effect suggests that Tke5 could be a valuable molecular tool for enhancing the biocontrol capabilities of *P. putida* in agricultural settings. By targeting multiple phytopathogens that threaten crops across various agricultural ecosystems, Tke5 represents a promising strategy for developing more effective and targeted biological control mechanisms.

## DISCUSSION

The identification and characterisation of novel protein toxins increase our molecular-level understanding of bacterial competition and the dynamics of microbial communities. This can be especially relevant in complex environments like the rhizosphere (surrounding plant roots) and phyllosphere (surrounding plant leaves), where rival bacteria compete to colonise plants [63]. Biocontrol agents in these complex environments, like *P. putida*, use the T6SS as a potent system to eliminate harmful phytopathogen competitors and protect crops from their infections. Here we describe Tke5, a novel potent protein toxin produced by *P. putida* with killing capabilities against the current most recalcitrant phytopathogens.

Tke5 is encoded within the K3-T6SS cluster and our bioinformatic analysis suggests it is a conserved T6SS component within the Pseudomonadota phylum (Fig. 1). Tke5 represents the first member of the BTH_I2691 family to be comprehensively studied at the molecular level. Another member of this family, VasX, was shown to be a *V. cholerae* T6SS-dependent virulence factor crucial to killing *Dictyostelium discoideum* [50] and later studies revealed it also functions as an antimicrobial effector, targeting the inner membrane of other *Vibrio* species leading to depolarisation and subsequent bacterial death [52]. Structural predictions of Tke5 suggest the presence of a MIX_III N-terminal (Fig. 1b), a well-established marker of T6SS effectors [12], which is also predicted in VasX. Additionally, Tke5 is predicted to contain multiple transmembrane domains (Supplementary Fig. 1a), a characteristic also present in VasX, which is associated with its membrane-targeting activity [52].

In the present study, we show that Tke5 exhibits potent antibacterial activity in growth inhibition assays using *P. putida* (Fig. 2b). Although we could not perform interbacterial competition assays due to the system’s inability to assemble under laboratory conditions, our results show that Tke5 is not merely bacteriostatic but actively kills target bacteria, mirroring the effects of a bactericidal antibiotic (Fig. 2c-d) without provoking cellular lysis. Flow cytometry data indicate that the bactericidal effect is due to cell depolarisation, which occurs without compromising cell permeability, suggesting that Tke5 permeabilises the inner membrane of the bacteria, while maintaining membrane integrity (Fig. 3).

To shed some light on the molecular mechanism underlying its cell depolarisation activity, we exploit electrophysiology using an *in vitro* model membrane (Supplementary Fig. 5). The observation of stable currents across the synthetic lipid bilayer upon Tke5 addition provides strong evidence for the formation of ion-selective pores (Fig. 4a-d). Further analysis of the RP under asymmetric salt concentrations (250/50 and 50/250 mM KCl in the cis/trans chambers) reveals that Tke5-induced pores allow the passive multi-ionic transport of ions, but with some preference for cations (K^+^) over anions (Cl^-^), as indicated by a calculated ion permeability ratio (P_+_/P_-_) ≍ 10. Furthermore, the RP is constant between traces of varying conductance levels (RP = −31 ±2 mV or +33 ±3 mV in a 250/50 or a 50/250 mM KCl gradient, respectively), which indicates Tke5 assembles a varying number of pores in each experimental replicate, but the structural characteristic of each pore is highly similar (Fig. 4e-g).

The capacity of Tke5 to permeate ions across the lipid bilayer is consistent with its cell depolarisation activity. Thus, we would expect that *in vivo*, Tke5 passively transports ions, with some preference for cations, from outside the cell, where they are found at higher concentrations, to inside the cell. This transport disrupts the delicate balance of ions across the membrane, which is crucial for maintaining ionic homeostasis and is essential for preserving the membrane potential and all the cellular processes that depend on it.

Several T6SS effectors have been identified that form ion-selective pores in the membranes of target cells, disrupting the membrane potential of the target cell and leading to cell death. Tse4 from *P. aeruginosa* was postulated to act by facilitating the formation of ion-selective membrane pores [64], and we have recently demonstrated that Tse4, indeed, promotes cell depolarisation in prey cells by incorporating into cellular membranes and forming multi-ionic channels [20]. Meanwhile, the C-terminal fragment of Tse5 (Tse5-CT) from the same strain is a PFT that assembles proteolipidic pores in the inner membrane of Gram-negative bacteria [19,65]. In this case, an ion permeability ratio value less than 6 in a 250/50 KCl gradient (P_+_/P_-_ ≤ 6) was measured using a model membrane with the same lipid composition (the *E. coli* polar lipid extract). Another example is Ssp6, found in *Serratia marcescens*, an antibacterial effector that mainly causes cell depolarisation [66]. For Ssp6, an RP = −26.5 mV was obtained in artificial membranes and non-symmetrical KCl conditions (510/210 mM KCl in *cis* and *trans* chambers, respectively).

Recently, a novel family of contact-dependent inhibition (CDI) effectors has been studied in *E. coli* [67]. This new family causes cell depolarisation, while minimal cell permeability was detected. Competition experiments demonstrate that these toxins are highly effective, with ∼82% of target cells showing depolarisation in representative assays. However, propidium iodide (PI) uptake, indicating loss of membrane integrity, was observed in only a minor fraction of cells. These results suggest these effectors might also be ionophoric protein toxins that assemble ion-selective membrane pores, causing cell depolarisation.

Unlike Tke5 and VasX, which are large proteins with approximately 1,000 amino acids, Tse5-CT, Ssp6, and the C-terminal fragments of CDI toxins represent a contrasting group of small toxins between ∼150-230 amino-acid residues. Despite significant differences in size, sequence, and origin, these toxins appear to converge on a similar mode of action, preferentially inserting into the inner membrane of intoxicated bacteria. This shared functional outcome underscores a fascinating case of convergent evolution in toxin mechanisms, likely driven by the same selective pressures to efficiently disrupt bacterial competitors.

Tke5 likely employs its N-terminal MIX domain for T6SS-dependent delivery into the target cell, while its C-terminal colicin-like domain is probably involved in causing cell depolarisation through ion channel formation. Our experimental data using synthetic lipid bilayers demonstrate that Tke5 forms stable, ion-selective pores with multi-ionic character and some preference for cations like K^+^, presumably destabilising the resting membrane potential of target bacteria. Notably, the measured ion conductance reveals that Tke5 inserts more efficiently under a 250/50 mM salt gradient, where the *cis* side of the bilayer, representing the periplasm, is more shielded than the *trans* side, which mimics the cytoplasm. This setup generates a negative membrane potential simulating physiological conditions, supporting the hypothesis that Tke5 inserts preferentially from the periplasm. Moreover, *in vivo* data show that Tke5 exhibits greater toxicity when directed to the Sec pathway compared to cytoplasmic expression (Fig. 2b), further supporting the periplasmic insertion model. A similar effect has been seen for Tse5, which senses the directionality of the membrane potential and can only release its encapsulated Tse5-CT fragment when introduced in the *cis* chamber under a 250/50 mM salt gradient, and not under a 50/250 mM salt gradient [19]. An even more dramatic effect is seen with Ssp6 from *Serratia*, which is not toxic when overproduced in the cytosol of *E. coli* and only forms ion channels when directed to the Sec translocon or delivered by the T6SS in a competition assay [66].

Tki5 expression protects *P. putida* from Tke5 toxicity (Fig. 2 and 3), confirming that Tke5/Tki5 functions as a genuine E/I pair. Tki5 is predicted to be a transmembrane protein (Supplementary Fig. 1c), which suggests it likely interacts with Tke5 at the membrane level to protect *P. putida* from self-intoxication and attack by sibling cells. A similar effect has been shown for VasX and its immunity protein TsiV2 [52]. However, the precise mechanism of this interaction remains to be elucidated.

In this study, we also highlight the ecological relevance of Tke5 by demonstrating its potent antibacterial activity against various plant pathogens. Expression of *tke5* significantly impaired growth in multiple phytopathogens, including *P. syringae, R. solanacearum, A. tumefaciens, D. dadantii, P. carotovorum, E. amylovora, X. campestris,* and *P. savastanoi*, compared to controls (Fig. 5). Control strains carrying empty vector or expressing non-toxic *msfGFP* grew normally with inducer (3-*m*Bz), confirming that growth inhibition stems directly from Tke5 activity. By effectively targeting and neutralising these economically important plant pathogens, Tke5 could play a crucial role in protecting crops and ensuring agricultural productivity. This would add a novel weapon to the *P. putida* arsenal after the already known effectors secreted by the K1-T6SS, such as the Tke2 nuclease [23] and the Tke7 colicin.

While T6SS is common in plant-associated bacteria [68], only a few T6SS effectors with biocontrol activity have been characterised. One example is the T6SS amidase Tae3^Pf^ from *P. fluorescens* MFE01, which protects against *Pectobacterium atrosepticum* [69]. Several beneficial bacteria employ T6SS for plant colonisation and pathogen defence, though their specific effectors remain uncharacterised. For instance, *Azospirillum brasilense* T6SS clusters provide antibacterial activity against multiple plant pathogens [70]. Similarly, *Pseudomonas chlororaphis* uses T6SSs to compete against pathogens like *A. tumefaciens* C58 and *P. syringae* pv. tomato DC3000 [71]. The F1- and F3-T6SS in *Pseudomonas ogarae* F113 are crucial for rhizosphere persistence, as demonstrated by the reduced fitness of T6SS-deficient mutants [72]. In *Pseudomonas fluorescens* Pf29Arp, T6SS genes show differential expression between healthy and pathogen-infected wheat roots, suggesting adaptation roles [73]. Even plant pathogens like *Acidovorax citrulli* seem to use T6SS to inhibit other pathogens, reducing rice bacterial blight symptoms caused by *X. oryzae* pv. oryzae [74]. In symbiotic bacteria, T6SS enhances host colonisation, as seen in the improved competitiveness of *Sinorhizobium freddii* in *G. max* cv Pekin nodules [75].

The characterisation of Tke5 as a novel, pore-forming, bactericidal toxin effector expands our understanding of the sophisticated weaponry employed by bacteria in interbacterial competition. Its activity against plant pathogens further suggests a key role in shaping microbial communities and influencing plant health. This knowledge can inform the development of novel biocontrol strategies and contribute to the ongoing efforts to reduce dependence on chemical pesticides.

## ACKNOWLEDGEMENTS

P.B. acknowledges the financial support received from the Spanish Minister of Science, Innovation and Universities (MICIU/AEI/10.13039/501100011033) through the *Ramón y Cajal* Program (RYC2019-026551-I, ESF Investing in your future), the research grant from the State Subprogram for Knowledge Generation PID2021-123000OB-I00 (ERDF/EU) and the research grant from the State Subprogram for Promotion of Research Consolidation CNS2022-135585 (European Union NextGenerationEU/PRTR).

D.A.-J. acknowledges support by the MICIU Contract PID2021-127816NB-I00 and the Basque Government’s Department of Education IT1745-22.

A.A. and M.Q.-M. acknowledge financial support by the Spanish Government MICIU/AEI/10.13039/501100011033/FEDER, UE (Project 2019-108434GB-I00 and Project PID2022-142795 NB-I00), Generalitat Valenciana (project CIGRIS/2021/021) and Universitat Jaume I (Project UJI-B2022-42). M.Q.-M acknowledges support from the Spanish Ministry of Science and Innovation (Project IJC2018-035283-I funded by MICIU/AEI/10.13039/501100011033) and Universitat Jaume I (project UJI-A2020-21).

We thank Cayo Ramos, Emilia López Solanilla, Francisco M. Cazorla, Ehr Min Lai, María Milagros López and Inmaculada Sampedro for their kind gifts of phytopathogenic strains.

## DATA AVAILABILITY STATEMENT

The data are available within the manuscript. The sequence of a newly generated vector available as a tool for the scientific community has been deposited into NCBI Genbank with accession number PQ628036 and into the SEVA repository https://seva-plasmids.com/.

## REFERENCES

1. Mondal AK, Verma P, Lata K, Singh M, Chatterjee S, Chattopadhyay K. Sequence Diversity in the Pore-Forming Motifs of the Membrane-Damaging Protein Toxins. Journal of Membrane Biology. Springer; 2020. pp. 469–478. doi:10.1007/s00232-020-00141-2

2. Carruthers VB. Apicomplexan Pore-Forming Toxins. Annu Rev Microbiol. 2024;78: 277–291. doi:10.1146/annurev-micro-041222

3. Peraro MD, Van Der Goot FG. Pore-forming toxins: Ancient, but never really out of fashion. Nature Reviews Microbiology. Nature Publishing Group; 2016. pp. 77–92. doi:10.1038/nrmicro.2015.3

4. Allsopp LP, Bernal P. Killing in the name of: T6SS structure and effector diversity. Microbiology (N Y). 2023;169: 1367. doi:10.1099/mic.0.001367

5. Rapisarda C, Cherrak Y, Kooger R, Schmidt V, Pellarin R, Logger L, et al. In situ and high-resolution cryo-EM structure of a bacterial type VI secretion system membrane complex. EMBO J. 2019;38: e100886. doi:10.15252/embj.2018100886

6. Cherrak Y, Rapisarda C, Pellarin R, Bouvier G, Bardiaux B, Allain F, et al. Biogenesis and structure of a type VI secretion baseplate. Nat Microbiol. 2018;3: 1404–1416. doi:10.1038/s41564-018-0260-1 PMID - 30323254

7. Zoued A, Durand E, Brunet YR, Spinelli S, Douzi B, Guzzo M, et al. Priming and polymerization of a bacterial contractile tail structure. Nature. 2016;531: 59–63. doi:10.1038/nature17182

8. Kudryashev M, Wang RYR, Brackmann M, Scherer S, Maier T, Baker D, et al. Structure of the Type VI secretion system contractile sheath. Cell. 2015;160: 952– 962. doi:10.1016/j.cell.2015.01.037

9. Basler M, Pilhofer M, Henderson GP, Jensen GJ, Mekalanos JJ. Type VI secretion requires a dynamic contractile phage tail-like structure. Nature. 2012;483: 182–186. doi:10.1038/nature10846

10. Bernal P, Furniss RCD, Fecht S, Leung RCY, Spiga L, Mavridou DAI, et al. A novel stabilization mechanism for the type VI secretion system sheath. Proc Natl Acad Sci U S A. 2021;118: e2008500118. doi:10.1073/pnas.2008500118

11. Santin YG, Doan T, Lebrun R, Espinosa L, Journet L, Cascales E. In vivo TssA proximity labelling during type VI secretion biogenesis reveals TagA as a protein that stops and holds the sheath. Nat Microbiol. 2018;3: 1304–1313. doi:10.1038/s41564-018-0234-3

12. Salomon D, Kinch LN, Trudgian DC, Guo X, Klimko JA, Grishin N V., et al. Marker for type VI secretion system effectors. Proc Natl Acad Sci U S A. 2014;111: 9271–9276. doi:10.1073/pnas.1406110111

13. Bernal P. T6SS-effector hunters uncover PIX: a novel delivery/marker domain. Trends Microbiol. 2024;32: 617–619. 10.1016/j.tim.2024.04.009

14. Kanarek K, Fridman CM, Bosis E, Salomon D. The RIX domain defines a class of polymorphic T6SS effectors and secreted adaptors. Nat Commun. 2023;14: 4983. doi:10.1038/s41467-023-40659-2

15. Jana B, Fridman CM, Bosis E, Salomon D. A modular effector with a DNase domain and a marker for T6SS substrates. Nat Commun. 2019;10: 3595. doi:10.1038/s41467-019-11546-6

16. Carobbi A, Leo K, Di Nepi S, Bosis E, Salomon D, Sessa G. PIX is an N-terminal delivery domain that defines a class of polymorphic T6SS effectors in *Enterobacterales*. Cell Rep. 2024;43: 114015. doi:10.1016/j.celrep.2024.114015

17. Fridman CM, Keppel K, Rudenko V, Altuna-Alvarez J, Albesa-Jové D, Bosis E, et al. A new class of type VI secretion system effectors can carry two toxic domains and are recognized through the WHIX motif for export. PLoS Biol. 2025;23. doi:10.1371/journal.pbio.3003053

18. González-Magaña A, Altuna J, Queralt-Martín M, Largo E, Velázquez C, Montánchez I, et al. The *P. aeruginosa* effector Tse5 forms membrane pores disrupting the membrane potential of intoxicated bacteria. Commun Biol. 2022;5: 1189. doi:10.1038/s42003-022-04140-y

19. González-Magaña A, Tascón I, Altuna-Alvarez J, Queralt-Martín M, Colautti J, Velázquez C, et al. Structural and functional insights into the delivery of a bacterial Rhs pore-forming toxin to the membrane. Nat Commun. 2023;14: 7808. doi:10.1038/s41467-023-43585-5

20. Rojas-Palomino J, Velazquez C, Altuna-Alvarez J, Gonzalez-Magana A, Zabala-Zearreta M, Muller M, et al. The Pseudomonas aeruginosa Tse4 toxin assembles ion-selective and voltage-sensitive ion channels to couple membrane depolarization with K+ efflux. PLoS Pathog. 2025;21. doi:10.1371/journal.ppat.1012981

21. Ruhe ZC, Low DA, Hayes CS. Polymorphic Toxins and Their Immunity Proteins: Diversity, Evolution, and Mechanisms of Delivery. Annu Rev Microbiol. 2020;74: 497–520. doi:10.1146/annurev-micro-020518

22. Weller DM. *Pseudomonas* Biocontrol Agents of Soilborne Pathogens: Looking Back Over 30 Years. Phytopathology. 2007;97: 250–256. doi:10.1094/phyto-97-2-0250 PMID - 18944383

23. Bernal P, Allsopp LP, Filloux A, Llamas MA. The *Pseudomonas putida* T6SS is a plant warden against phytopathogens. ISME Journal. 2017;11: 972–987. doi:10.1038/ismej.2016.169

24. Bernal P, Civantos C, Pacheco-Sánchez D, Quesada JM, Filloux A, Llamas MA. Transcriptional organization and regulation of the *Pseudomonas putida* K1 type VI secretion system gene cluster. Microbiology (N Y). 2023;169: 001295. doi:10.1099/mic.0.001295

25. Sambrook J, Fritsch EF, Maniatis T. Molecular Cloning: A Laboratory Manual. 2nd edition. Cold Spring Harbor Laboratory, editor. NY: Molecular Cloning: A Laboratory Manual; 1989.

26. Choi K-H, Kumar A, Schweizer HP. A 10-min method for preparation of highly electrocompetent *Pseudomonas aeruginosa* cells: application for DNA fragment transfer between chromosomes and plasmid transformation. J Microbiol Methods. 2006;64: 391–397. doi:10.1016/j.mimet.2005.06.001

27. Ramos-Gonzalez MI, Duque E, Ramos JL. Conjugational transfer of recombinant DNA in cultures and in soils: host range of *Pseudomonas putida* TOL plasmids. Appl Environ Microbiol. 1991;57: 3020–3027.

28. Lei S-P, Lin H-C, Wang S-S, Callaway J, Wilcox G. Characterization of the *Erwinia carotovora pelB* Gene and Its Product Pectate Lyase. J Bacteriol. 1987;169: 4379– 4383. Available: https://journals.asm.org/journal/jb

29. Tsirigotaki A, De Geyter J, Šoštarić N, Economou A, Karamanou S. Protein export through the bacterial Sec pathway. Nat Rev Microbiol. 2017;15: 21–36. doi:10.1038/nrmicro.2016.161

30. Calles B, Goñi-Moreno Á, Lorenzo V. Digitalizing heterologous gene expression in Gram-negative bacteria with a portable ON/OFF module. Mol Syst Biol. 2019;15: e8777. doi:10.15252/msb.20188777

31. Silva-Rocha R, Martínez-García E, Calles B, Chavarría M, Arce-Rodríguez A, Heras A de las, et al. The Standard European Vector Architecture (SEVA): a coherent platform for the analysis and deployment of complex prokaryotic phenotypes. Nucleic Acids Res. 2013;41: D666–D675. doi:10.1093/nar/gks1119 PMID - 23180763

32. Martínez-García E, Fraile S, Algar E, Aparicio T, Velázquez E, Calles B, et al. SEVA 4.0: an update of the Standard European Vector Architecture database for advanced analysis and programming of bacterial phenotypes. Nucleic Acids Res. 2023;51: D1558–D1567. doi:10.1093/nar/gkac1059

33. Nikel PI, Benedetti I, Wirth NT, de Lorenzo V, Calles B. Standardization of regulatory nodes for engineering heterologous gene expression: a feasibility study. Microb Biotechnol. 2022;15: 2250–2265. doi:10.1111/1751-7915.14063

34. Winsor GL, Griffiths EJ, Lo R, Dhillon BK, Shay JA, Brinkman FSL. Enhanced annotations and features for comparing thousands of *Pseudomonas* genomes in the *Pseudomonas* genome database. Nucleic Acids Res. 2016;44: D646–D653. doi:10.1093/nar/gkv1227

35. Boratyn GM, Camacho C, Cooper PS, Coulouris G, Fong A, Ma N, et al. BLAST: a more efficient report with usability improvements. Nucleic Acids Res. 2013;41: W29–W33. doi:10.1093/nar/gkt282

36. Letunic I, Doerks T, Bork P. SMART: Recent updates, new developments and status in 2015. Nucleic Acids Res. 2015;43: D257–D260. doi:10.1093/nar/gku949

37. Finn RD, Coggill P, Eberhardt RY, Eddy SR, Mistry J, Mitchell AL, et al. The Pfam protein families database: towards a more sustainable future. Nucleic Acids Res. 2016;44: D279–D285. doi:10.1093/nar/gkv1344 PMID - 26673716

38. Wang J, Chitsaz F, Derbyshire MK, Gonzales NR, Gwadz M, Lu S, et al. The conserved domain database in 2023. Nucleic Acids Res. 2023;51: D384–D388. doi:10.1093/NAR/GKAC1096

39. Krogh A, Larsson B, Von Heijne G, Sonnhammer ELL. Predicting transmembrane protein topology with a hidden markov model: application to complete genomes. J Mol Biol. 2001;305: 567–580. doi:10.1006/JMBI.2000.4315

40. Tamura K, Stecher G, Kumar S. MEGA11: Molecular Evolutionary Genetics Analysis Version 11. Mol Biol Evol. 2021;38: 3022–3027. doi:10.1093/MOLBEV/MSAB120

41. Jones DT, Taylor WR, Thornton JM. The rapid generation of mutation data matrices from protein sequences. Comput Appl Biosci. 1992;8: 275–282. doi:10.1093/BIOINFORMATICS/8.3.275

42. Letunic I, Bork P. Interactive Tree Of Life (iTOL) v4: recent updates and new developments. Nucleic Acids Res. 2019;47: W256–W259. doi:10.1093/NAR/GKZ239

43. van Kempen M, Kim SS, Tumescheit C, Mirdita M, Lee J, Gilchrist CLM, et al. Fast and accurate protein structure search with Foldseek. Nat Biotechnol. 2024;42: 243–246. doi:10.1038/s41587-023-01773-0

44. Madeira F, Madhusoodanan N, Lee J, Eusebi A, Niewielska A, Tivey ARN, et al. The EMBL-EBI Job Dispatcher sequence analysis tools framework in 2024. Nucleic Acids Res. 2024;52: W521–W525. doi:10.1093/nar/gkae241

45. Meng EC, Goddard TD, Pettersen EF, Couch GS, Pearson ZJ, Morris JH, et al. UCSF ChimeraX: Tools for structure building and analysis. Protein Science. 2023;32: e4792. doi:10.1002/pro.4792

46. Montal M, Mueller P. Formation of Bimolecular Membranes from Lipid Monolayers and a Study of Their Electrical Properties. Proceedings of the National Academy of Sciences. 1972;69: 3561–3566. doi:10.1073/PNAS.69.12.3561

47. Morein S, Andersson A-S, Rilfors L, Ran Lindblom G. Wild-type *Escherichia coli* Cells Regulate the Membrane Lipid Composition in a “Window” between Gel and Non-lamellar Structures*. J Biol Chem. 1996;271: 6801–6809.

48. Alcaraz A, Nestorovich EM, López ML, García-Giménez E, Bezrukov SM, Aguilella VM. Diffusion, exclusion, and specific binding in a large channel: A study of OmpF selectivity inversion. Biophys J. 2009;96: 56–66. doi:10.1016/j.bpj.2008.09.024

49. Effect THE, Sodium OF, Activity E, The OF. Laboratory, University. 1948; 37–77.

50. Miyata ST, Kitaoka M, Brooks TM, McAuley SB, Pukatzki S. *Vibrio cholerae* Requires the Type VI Secretion System Virulence Factor VasX To Kill *Dictyostelium discoideum*. Infect Immun. 2011;79: 2941–2949. doi:10.1128/iai.01266-10 PMID - 21555399

51. Dong TG, Ho BT, Yoder-Himes DR, Mekalanos JJ. Identification of T6SS-dependent effector and immunity proteins by Tn-seq in *Vibrio cholerae*. Proc Natl Acad Sci U S A. 2013;110: 2623–2628. doi:10.1073/pnas.1222783110

52. Miyata ST, Unterweger D, Rudko SP, Pukatzki S. Dual Expression Profile of Type VI Secretion System Immunity Genes Protects Pandemic *Vibrio cholerae*. PLoS Pathog. 2013;9: e1003752. doi:10.1371/journal.ppat.1003752

53. Sockolosky JT, Szoka FC. Periplasmic production via the pET expression system of soluble, bioactive human growth hormone. Protein Expr Purif. 2013;87: 129–135. doi:10.1016/J.PEP.2012.11.002

54. Buller GM, Bradford JA, Liu J, Godfrey WL. Novel Reagents for the Addition of Viability Measurements to Immunostaining Using Flow Cytometry. Blood. 2006;108: 3879–3879. doi:10.1182/BLOOD.V108.11.3879.3879

55. Winkel JD te, Gray DA, Seistrup KH, Hamoen LW, Strahl H. Analysis of antimicrobial-triggered membrane depolarization using voltage sensitive dyes. Front Cell Dev Biol. 2016;4: 29. doi:10.3389/FCELL.2016.00029/BIBTEX

56. LaCourse KD, Peterson SB, Kulasekara HD, Radey MC, Kim J, Mougous JD. Conditional toxicity and synergy drive diversity among antibacterial effectors. Nat Microbiol. 2018;3: 440–446. doi:10.1038/s41564-018-0113-y

57. Aguilella VM, Queralt-Martín M, Aguilella-Arzo M, Alcaraz A. Insights on the permeability of wide protein channels: measurement and interpretation of ion selectivity. Integrative Biology. 2011;3: 159–172. doi:10.1039/C0IB00048E

58. Alcaraz A, Nestorovich EM, López ML, García-Giménez E, Bezrukov SM, Aguilella VM. Diffusion, exclusion, and specific binding in a large channel: a study of OmpF selectivity inversion. Biophys J. 2009;96: 56–66. doi:10.1016/J.BPJ.2008.09.024

59. Hille B. Ion Channels of Excitable Membranes. Third Edition. Sunderland, MA, MA: Sinauer Associates, Inc; 2001.

60. Park HB, Kamcev J, Robeson LM, Elimelech M, Freeman BD. Maximizing the right stuff: The trade-off between membrane permeability and selectivity. Science (1979). 2017;356: 1138–1148. doi:10.1126/SCIENCE.AAB0530/ASSET/1FAB46CE-E045-4C54-80A4-6FD3685D9AEA/ASSETS/GRAPHIC/356_AAB0530_F5.JPEG

61. Hodgkin AL, Katz B. The effect of sodium ions on the electrical activity of the giant axon of the squid. J Physiol. 1949;108: 37–77. doi:10.1113/JPHYSIOL.1949.SP004310

62. Mansfield J, Genin S, Magori S, Citovsky V, Sriariyanum M, Ronald P, et al. Top 10 plant pathogenic bacteria in molecular plant pathology. Molecular Plant Pathology. 2012. pp. 614–629. doi:10.1111/j.1364-3703.2012.00804.x

63. Vázquez D, Civantos C, Durán-Wendt D, Ruiz A, Rivilla R, Martín M, et al. The *Pseudomonas putida* Type VI Secretion Systems Shape the Tomato Rhizosphere Microbiota. BioRxiv. 2025. doi:10.1101/2025.08.14.670259

64. LaCourse KD, Peterson SB, Kulasekara HD, Radey MC, Kim J, Mougous JD. Conditional toxicity and synergy drive diversity among antibacterial effectors. Nat Microbiol. 2018;3: 440–446. doi:10.1038/s41564-018-0113-y

65. Rojas-Palomino J, Altuna-Alvarez J, González-Magaña A, Queralt-Martín M, Albesa-Jové D, Alcaraz A. Electrophysiological dissection of the ion channel activity of the Pseudomonas aeruginosa ionophore protein toxin Tse5. Chem Phys Lipids. 2025;267. doi:10.1016/j.chemphyslip.2025.105472

66. Mariano G, Trunk K, Williams DJ, Monlezun L, Strahl H, Pitt SJ, et al. A family of Type VI secretion system effector proteins that form ion-selective pores. Nat Commun. 2019;10: 5484. doi:10.1038/s41467-019-13439-0

67. Halvorsen TM, Schroeder KA, Jones AM, Hammarlöf D, Low DA, Koskiniemi S, et al. Contact-dependent growth inhibition (CDI) systems deploy a large family of polymorphic ionophoric toxins for inter-bacterial competition. Tan S, editor. PLoS Genet. 2024;20: e1011494. doi:10.1371/journal.pgen.1011494

68. Bernal P, Llamas MA, Filloux A. Type VI secretion systems in plant-associated bacteria. Environ Microbiol. 2018;20: 1–15. doi:10.1111/1462-2920.13956

69. Bourigault Y, Dupont CA, Desjardins JB, Doan T, Bouteiller M, Le Guenno H, et al. *Pseudomonas fluorescens* MFE01 delivers a putative type VI secretion amidase that confers biocontrol against the soft-rot pathogen *Pectobacterium atrosepticum*. Environ Microbiol. 2023;25: 2564–2579. doi:10.1111/1462-2920.16492

70. Cassan FD, Coniglio A, Amavizca E, Maroniche G, Cascales E, Bashan Y, et al. The *Azospirillum brasilense* Type VI secretion system promotes cell aggregation, biocontrol protection against phytopathogens and attachment to the microalgae *Chlorella sorokiniana*. Environ Microbiol. 2021;23: 6257–6274. doi:10.1111/1462-2920.15749

71. Boak EN, Kirolos S, Pan H, Pierson LS, Pierson EA. The Type VI Secretion Systems in Plant-Beneficial Bacteria Modulate Prokaryotic and Eukaryotic Interactions in the Rhizosphere. Front Microbiol. 2022;13: 843092. doi:10.3389/fmicb.2022.843092

72. Durán D, Bernal P, Vazquez-Arias D, Blanco-Romero E, Garrido-Sanz D, Redondo-Nieto M, et al. *Pseudomonas fluorescens* F113 type VI secretion systems mediate bacterial killing and adaption to the rhizosphere microbiome. Sci Rep. 2021;11: 5772. doi:10.1038/s41598-021-85218-1

73. Marchi M, Boutin M, Gazengel K, Rispe C, Gauthier J, Guillerm-Erckelboudt A, et al. Genomic analysis of the biocontrol strain *Pseudomonas fluorescens* Pf29Arp with evidence of T3SS and T6SS gene expression on plant roots. Environ Microbiol Rep. 2013;5: 393–403. doi:10.1111/1758-2229.12048 PMID - 23754720

74. Kan Y, Zhang Y, Lin W, Dong T. Differential plant cell responses to *Acidovorax citrulli* T3SS and T6SS reveal an effective strategy for controlling plant-associated pathogens. mBio. 2023;14: e0045923. doi:10.1128/mbio.00459-23

75. Reyes-Pérez PJ, Jiménez-Guerrero I, Sánchez-Reina A, Civantos C, Castro NM, Ollero FJ, et al. The Type VI Secretion System of *Sinorhizobium fredii* USDA257 Is Required for Successful Nodulation With *Glycine max* cv Pekin. Microb Biotechnol. 2025;18: e70112. doi:10.1111/1751-7915.70112

