## Supplementary Data and Datasets for "Tke5 is a novel *Pseudomonas putida* toxin that depolarises membranes killing plant pathogens"

**This file includes:**

Figures S1 to S5

Tables S1 to S3

Legends for Dataset S1 and S2

Supplementary references

SUPPLEMENTARY INFORMATION FIGURES

Figure S1

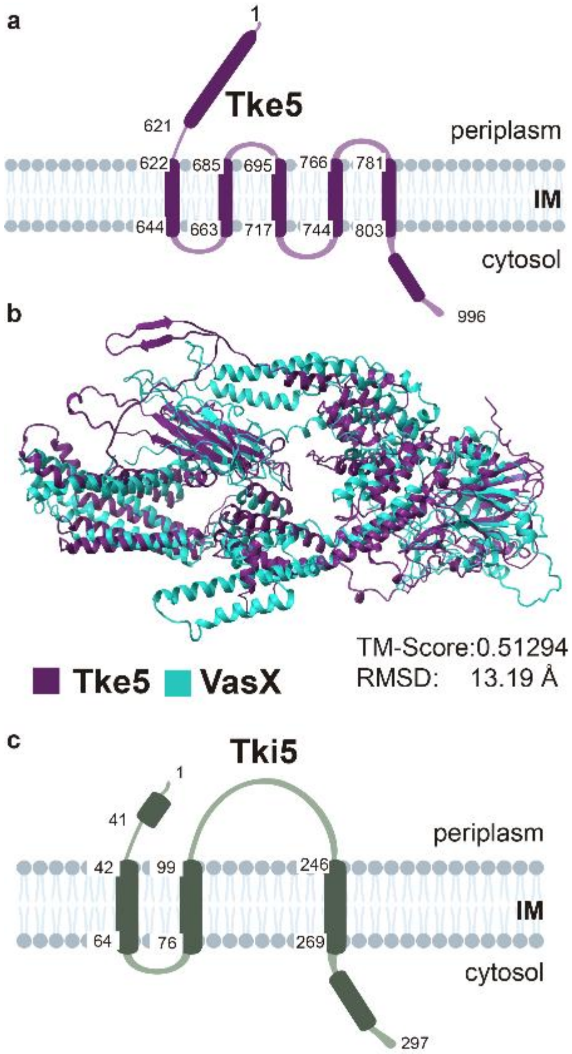


**Supplementary Figure 1 | Tke5 and Tki5 *in silico* studies a)** Topology prediction of Tke5 effector from *P. putida* T6SS. Tke5 transmembrane domain prediction was generated using TMHMM-2.0. **b)** Structural alignment of the AlphaFold Tke5 (purple) and VasX (cyan) models using Foldseek. **c)** Topology prediction of Tki5 immunity protein from *P. putida* T6SS. Tke5 transmembrane domain prediction was generated using TMHMM-2.0.

#### Figure S2

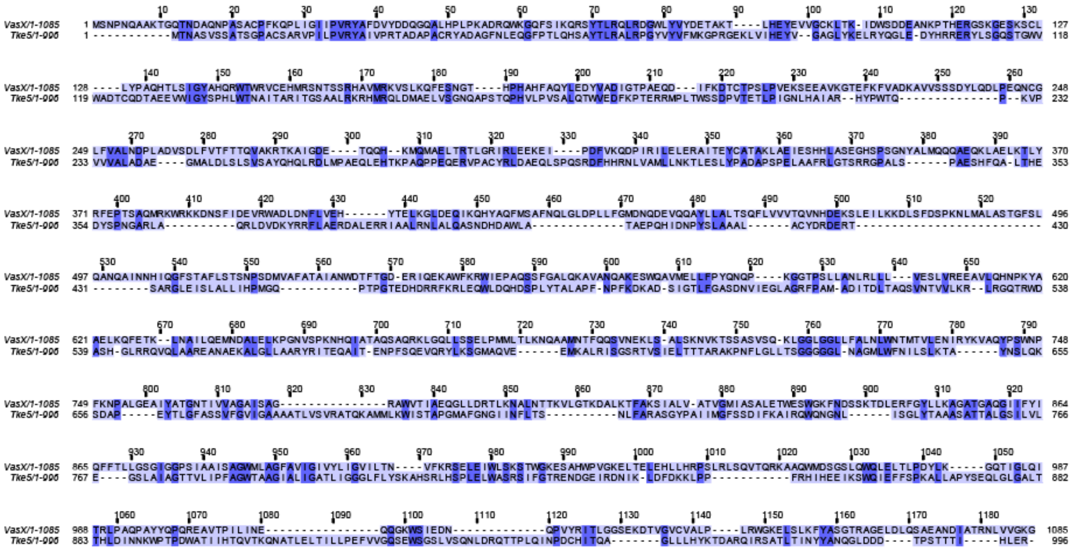

Percent Identity Matrix - created by Clustal2.1

| | | |
|---|---|---|
| 1: VasX | 100.00 | 21.36 |
| 2: Tke5 | 21.36 | 100.00 |

**Supplementary Figure 2 |** Multiple Sequence Alignment (MSA) of Tke5 and VasX sequences using Clustal Omega and displayed by Jalview coloured by percentage of identity. The percent identity matrix created by Clustal 2.1 is included in the bottom part of the figure.

#### Figure S3

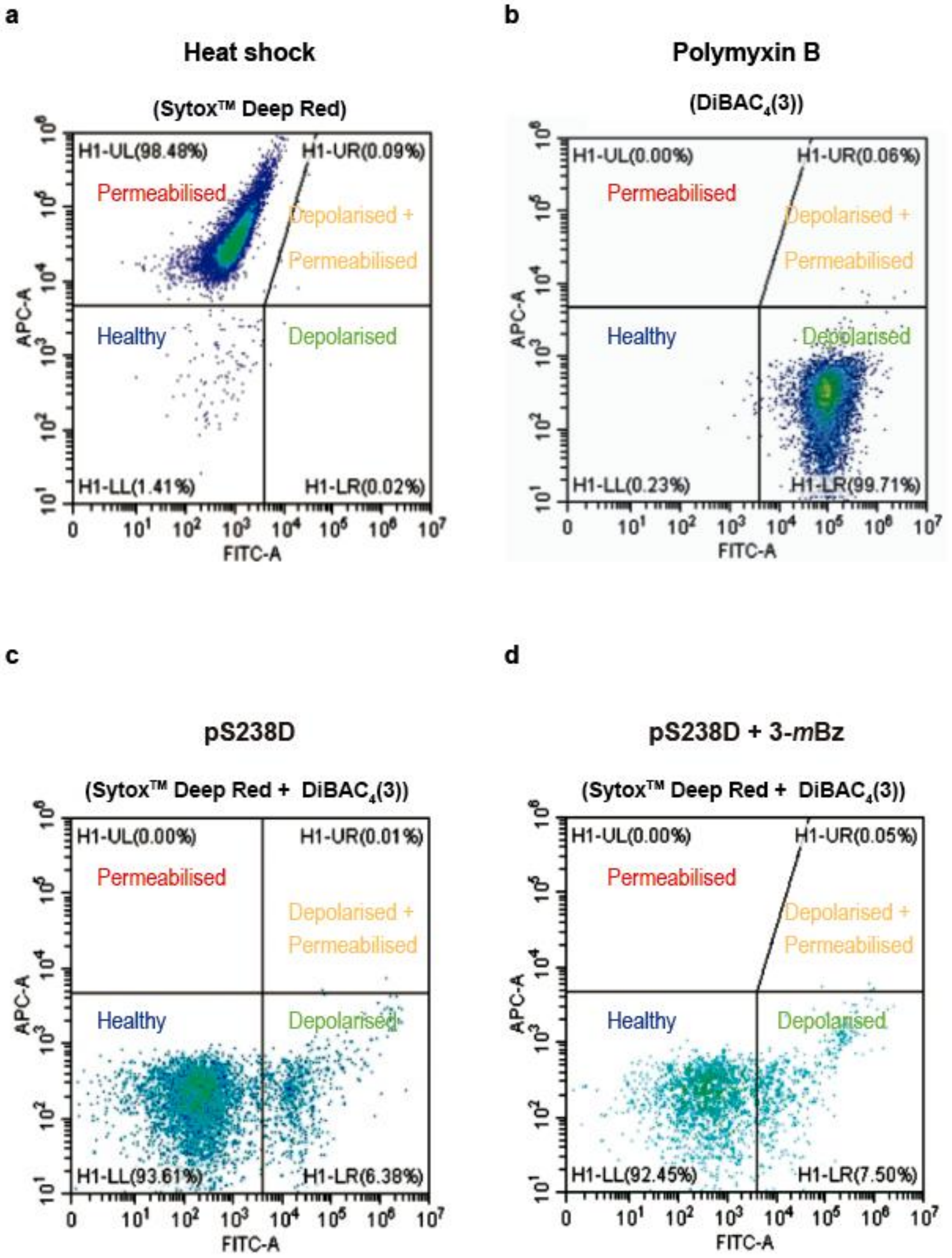
a
Heat shock
(Sytox™ Deep Red)
H1-UL(98.48%)
H1-UR(0.09%)
Permeabilised
Depolarised +
Permeabilised
Healthy
Depolarised
H1-LL(1.41%)
H1-LR(0.02%)
APC-A
FITC-A
b
Polymyxin B
(DiBAC4(3))
H1-UL(0.00%)
H1-UR(0.06%)
Permeabilised
Depolarised +
Permeabilised
Healthy
Depolarised
H1-LL(0.23%)
H1-LR(99.71%)
APC-A
FITC-A
c
pS238D
(Sytox™ Deep Red + DiBAC4(3))
H1-UL(0.00%)
H1-UR(0.01%)
Permeabilised
Depolarised +
Permeabilised
Healthy
Depolarised
H1-LL(93.61%)
H1-LR(6.38%)
APC-A
FITC-A
d
pS238D + 3-mBz
(Sytox™ Deep Red + DiBAC4(3))
H1-UL(0.00%)
H1-UR(0.05%)
Permeabilised
Depolarised +
Permeabilised
Healthy
Depolarised
H1-LL(92.45%)
H1-LR(7.50%)
APC-A
FITC-A

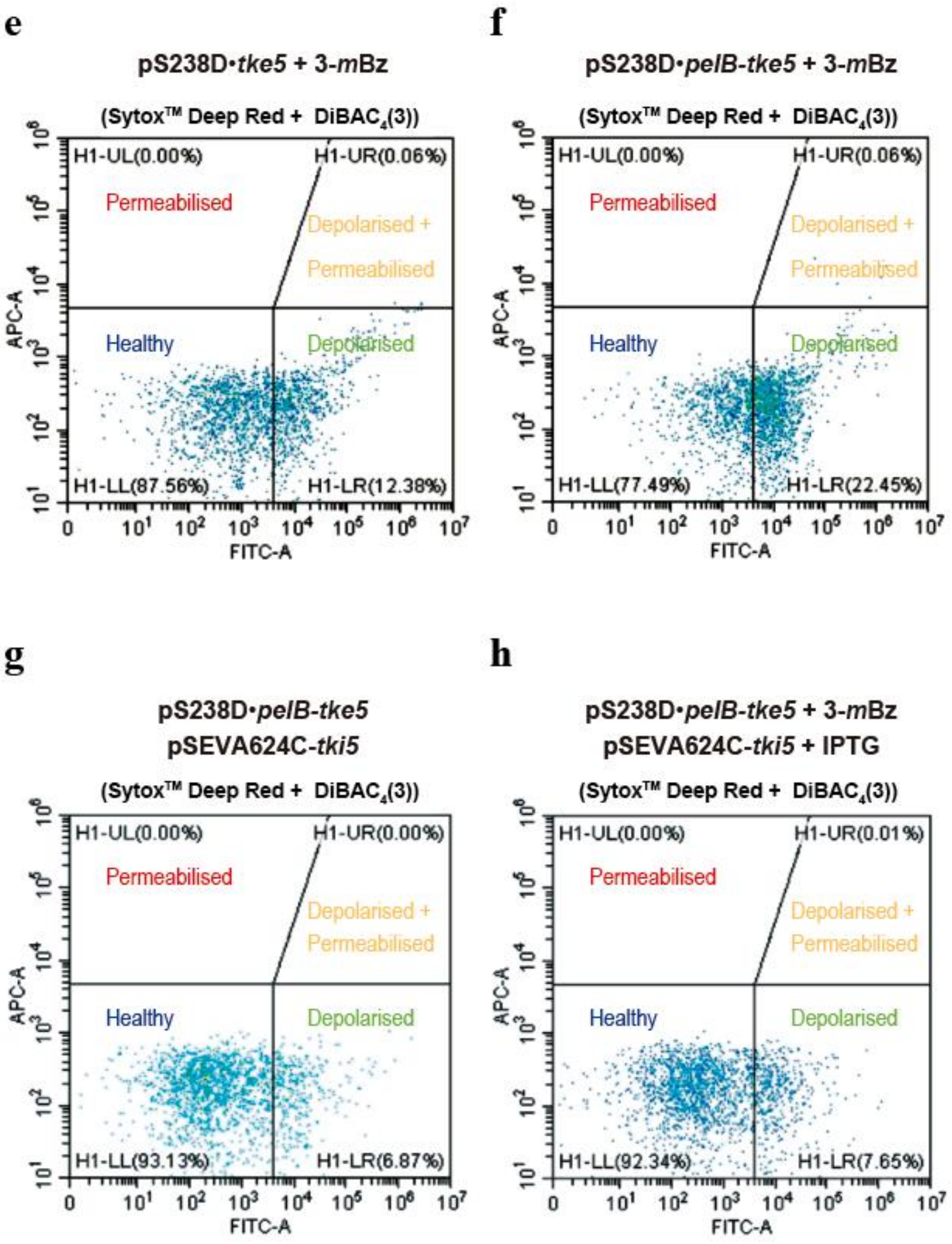


**Supplementary Figure 3 | Depolarisation and permeabilisation controls analysed by flow cytometry. a)** *E. coli* DH5α cells upon heat shock treatment at 85 ºC for 2 min and dyed with Sytox$^{TM}$ Deep Red, which emits red fluorescence (APC-A channel) as a marker for permeabilised cells. **b)** *E. coli* DH5α cells upon treatment with polymyxin B sulfate (100 μg/mL) and dyed with DiBAC$_4$(3), which emits green fluorescence (FITC-A channel) as a marker for depolarised cells. **c)** *E. coli* DH5α cells harbouring the empty pS238D vector and dyed with Sytox$^{TM}$ Deep Red and DiBAC$_4$(3) to determine the healthy bacteria population. **d)** *E. coli* DH5α cells harbouring the empty pS238D vector and treated with the inductor 3-*m*Bz before dying with Sytox$^{TM}$ Deep Red and DiBAC$_4$(3) to determine any putative side effect of this molecule in permeabilisation and/or depolarisation. **e)** *E. coli* DH5α cells expressing Tke5 upon induction with 3-*m*Bz (pS238D•*tke5*) and stained with DiBAC$_4$(3) and Sytox$^{TM}$ Deep Red. **f)** *E. coli* DH5α cells expressing PelB-Tke5 upon induction with 3-*m*Bz (pS238D•*pelB-tke5*) and stained with DiBAC$_4$(3) and Sytox$^{TM}$ Deep Red. **g)** *E. coli* DH5α cells harboruing pS238D•*pelB-tke5* and pSEVA624C-*tki5* and stained with DiBAC$_4$(3) and Sytox$^{TM}$ Deep Red as a control. **h)** *E. coli* DH5α cells coexpressing PelB-Tke5 and Tki5 (pS238D•*pelB-tke5* and pSEVA624C-*tki5*) upon IPTG and 3-*m*Bz induction and stained with DiBAC$_4$(3) and Sytox$^{TM}$ Deep Red.

**Figure S4**

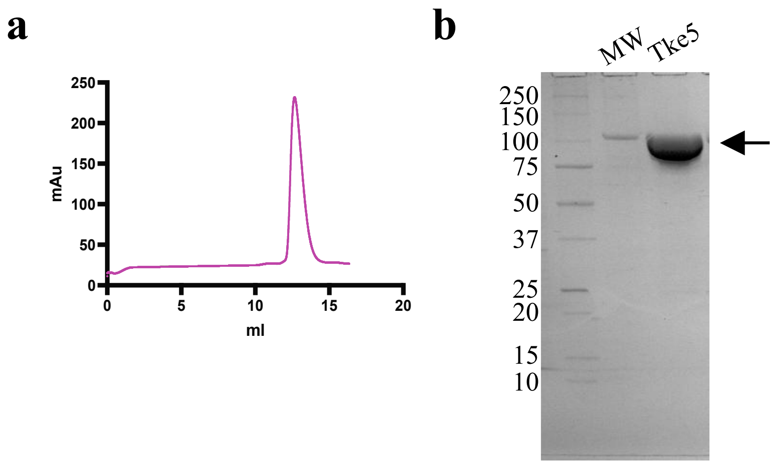


**Supplementary Figure 4 | Expression and purification of *P. putida* Tke5 effector.** **A)** Chromatogram of a size exclusion chromatography (Superdex™ 200 Increase 10/300 GL) of Tke5 shows only one peak that approximates the molecular weight of its monomer (112.6 kDa). **b)** Sodium dodecyl sulphate – polyacrylamide gel electrophoresis (SDS-PAGE) of purified Tke5 shows a unique band slightly higher than 100 kDa.

**Figure S5**

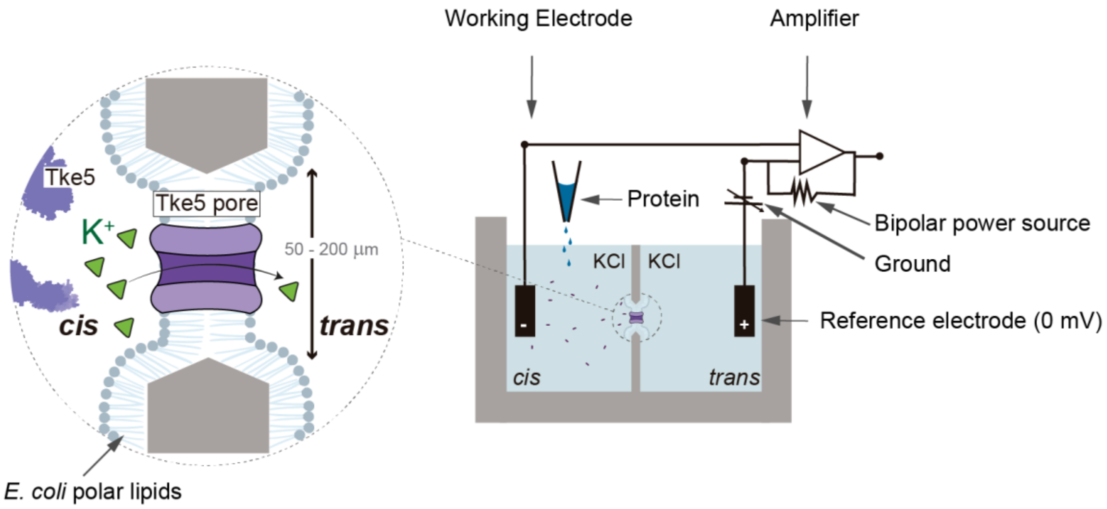


**Supplementary Figure 5 |** Schematic of a Montal-Mueller setup to form a synthetic membrane from an *E. coli* polar lipid extract.

#### SUPPLEMENTARY INFORMATION TABLES

**Table S1.** Bacterial strains used in this study. The antibiotic resistance markers are identified as follows: Amp, ampicillin; Km, kanamycin; Gm, gentamicin, Sm, streptomycin, Nal, nalidixic acid; Pip, piperacillin and Rif, rifampicin.

| Name | Description | Source |
|---|---|---|
| ***Escherichia coli*** | | |
| DH5α | F$^-$ *end*A1 *gln*V44 *thi*-1 *rec*A1 *rel*A1 *gyr*A96 *deo*R *nup*G *pur*B20 φ80d*lacZ*ΔM15 Δ(*lac*ZYA-*arg*F*)*U169 *hsd*R17(r$_K^-$m$_K^+$) λ$^-$, Nal$^R$ | (Hanahan, 1985) |
| CC118λpir | *ara*D Δ(*ara*, *leu*) Δ*lac*Z74 *pho*A20 *gal*K *thi*-1 *rsp*E *rpo*B *arg*E *rec*A1 λ*pir*, Rif$^R$ | (Herrero *et al.*, 1990) |
| HB101 | *supE44 hsdS20 recA13 ara-14 proA2 lacY1 galK2 rpsL20 xyl-5 mtl-1*, Sm$^R$ | (Boyer and Roulland-Dussoix, 1969) |
| BL21(DE3) | F$^-$*, ompT, hsdSB(rB–mB–), gal, Ion, λ(DE3 [lacI lacUV5-T7p07 ind1 sam7 nin5]) [malB+]*K$^-$*12(λS), dcm.* | (Studier and Moffatt, 1986) |
| ***Pseudomonas putida*** | | |
| KT2440R | Wild type strain, Rif$^R$ | (Espinosa-Urgel *et al.*, 2000) |
| ***Plant pathogens*** | | |
| *Pseudomonas syringae* pv. syringae B728a | | (Feil *et al.*, 2005) |
| *Ralstonia solanacearum* NCPPB 1493 (CECT 125) | | CECT (Spanish Collection of Type Cultures) |
| *Agrobacterium tumefaciens* C58 | | Erh Min Lai collection (Academia Sinica, Taiwan) |
| *Dickeya dadantii* 3937 (*Erwinia chrysanthemi* 3937) | | (Kotoujansky *et al.*, 1985) |
| *Pectobacterium carotovorum* subsp. carotovorum SCRI 194 | | María Milagros Lopez collection (IVIA, Spain) |
| *Erwinia amylovora* NCPPB 595 (CECT 222) | | CECT (Spanish Collection of Type Cultures) |
| *Xanthomonas campestris* pv vesicatoria NCPPB 195 (CECT 792) | | CECT (Spanish Collection of Type Cultures) |
| *Pseudomonas savastanoi* pv. savastanoi NCPPB 3335 | | (Pérez-Martínez *et al.*, 2007) |

**Table S2.** Plasmids used in this study. The antibiotic resistance markers are identified as follows: Amp, ampicillin; Km, kanamycin; Gm, gentamicin, Sm, streptomycin, Cm, chloramphenicol, Pip, piperacillin and Rif, rifampicin.

| Name | Description | Source |
|---|---|---|
| pJET1.2/blunt | Cloning vector, ColE1 *ori*, $Amp^R$ | Thermo Scientific |
| pRK600 | Helper plasmid, ColE1 *ori, mob*RK2, *tra*RK2, $Cm^R$ | (Kessler *et al.*, 1992) |
| pS238D•M | Tightly regulated expression vector, pBBR1 *ori*, *xylS*-$P_m$, *msf*•GFP, $Km^R$ | (Calles *et al.*, 2019) |
| pS238D•*tke5* | *tke5* encoding Tke5 cloned into pS238D•M, pBBR1 *ori*, *xylS*-$P_m$, $Km^R$ | This study |
| pS238D•*pelB-tke5* | *tke5* encoding Tke5 with an N-terminal PelB leader sequence cloned into pS238D•M, pBBR1 *ori*, *xylS*-$P_m$, $Km^R$ | This study |
| pSEVA621 | Expression vector, RK2 *ori*, $Gm^R$ | (Silva-Rocha *et al.*, 2013) |
| pSEVA234C | Expression vector, pBBR1 *ori*, *lacI*$^q$-$P_{trc}$, $Km^R$ | (Nikel *et al.*, 2022) |
| pSEVA624C | Expression vector derivate from pSEVA621 where the *lacI*$^q$ is divergent to the $P_{trc}$ (*lacI*$^q$-$P_{trc}$ region from pSEVA234C), RK2 *ori*, $Gm^R$ | This study |
| pSEVA424 | Expression vector, RK2 *ori*, *lacI*$^q$-$P_{trc}$, $Sm^R$ | (Silva-Rocha *et al.*, 2013; Martínez-García *et al.*, 2023) |
| pSEVA424•*tki5* | *tki5* encoding Tki5 preceded by an artificial RBS and cloned into pSEVA424, RK2 *ori*, *lacI*$^q$-$P_{trc}$, $Sm^R$ | This study |
| pSEVA624C•*tki5* | *tki5* encoding Tki5 from pSEVA424-*tki5* preceded by an artificial RBS and *sub*cloned into pSEVA624C, RK2 *ori*, *lacI*$^q$-$P_{trc}$, $Gm^R$ | This study |
| pCOLADuet™-1 | Expression vector with MCS-1 and MCS-2 for coexpression of two target genes, ColA *ori*, *lacI*$^q$-$P_{T7/lacO}$, $Km^R$ | Novagen |
| pCOLADuet-1•*9xhis-tke5* | *tke5* encoding Tke5 with an N-terminal 9xHis tag and TEV protease cleavage site cloned into *mcs-2* of pCOLADuet™-1, ColA *ori*, *lacI*$^q$-$P_{T7/lacO}$, $Km^R$ | This study |

**Table S3.** Oligonucleotide primers used in this study. The "Brief description" column provides basic information on the primer design (restriction enzyme used for cloning, encoded protein, forward or reverse orientation of the primer (F or R); OP stands for Overlapping PCR, after the / symbol it is indicated the vector where the PCR product has been cloned.

| Number | Brief description | Sequence (5´-3´) | PB code |
|---|---|---|---|
| P1 | NheI.*tke5*.F/ pS238D•M | aattaaGCTAGCactaacgcctccgtaagcag | PB0644 |
| P2 | BamHI.*tke5*.R/ pS238D•M | ggccatGGATCCgtttagcgttccagatgaatagt gg | PB0647 |
| P3 | NheI.*pelBtke5*.F /pS238D•M | aattaaGCTAGC*aagtacctgctgccgaccgcc gccgccggcctgctgctgctggccgcccagccggcc atggcc*actaacgcctccgtaagcag | PB0645 |
| P4 | S238D-F | tgctgcaactctctcaggg | PB0627 |
| P5 | S238D-R | gggttttcccagtcacgac | PB0003 |
| P6 | SEVA624C-F | agggcggcggatttgtcc | PB0742 |
| P7 | SEVA624C-R | gcggcaaccgagcgttc | PB0743 |
| P8 | SEVA-F | agcggataacaatttcacacagga | PB101 |
| P9 | SEVA-R | cgccagggttttcccagtcacgac | PB100 |

### DATASETS

#### Dataset 1

| Index | Query | Status | Score | Sequences | Taxonomy | Taxid | Accessions | Domain Names |
|---|---|---|---|---|---|---|---|---|
| 1 | ="p469444" | done | 1 | 7417 | cellular organisms | 131567 | NF041559 | BTH_I2691_fam |
| 1 | ="p469444" | done | 1 | 2958 | Bacteria | 2 | cl41762~NF041559 | MIX~BTH_I2691_fam |
| 1 | ="p469444" | done | 1 | 856 | Bacteria | 2 | NF041559~cl41762 | BTH_I2691_fam~MIX |
| 1 | ="p469444" | done | 1 | 100 | Pseudomonas | 286 | cl41762~NF041559~COG4942 | MIX~BTH_I2691_fam~EnvC |
| 1 | ="p469444" | done | 1 | 58 | Bacteria | 2 | pfam20249~NF041559 | VasX_N~BTH_I2691_fam |
| 1 | ="p469444" | done | 1 | 35 | Gammaproteobacteria | 1236 | cl41762~COG4942~NF041559 | MIX~EnvC~BTH_I2691_fam |
| 1 | ="p469444" | done | 1 | 18 | Gammaproteobacteria | 1236 | cl41762~COG4372~NF041559 | MIX~COG4372~BTH_I2691_fam |
| 1 | ="p469444" | done | 1 | 15 | Gammaproteobacteria | 1236 | cl41762~COG3321~NF041559 | MIX~PksD~BTH_I2691_fam |
| 1 | ="p469444" | done | 1 | 13 | Gammaproteobacteria | 1236 | cl41762~TIGR02168~NF041559 | MIX~SMC_prok_B~BTH_I2691_fam |
| 1 | ="p469444" | done | 1 | 10 | Gammaproteobacteria | 1236 | COG1388~cl41762~NF041559 | LysM~MIX~BTH_I2691_fam |
| 1 | ="p469444" | done | 1 | 10 | Gammaproteobacteria | 1236 | cl46249~cl41762~NF041559 | LysM~MIX~BTH_I2691_fam |
| 1 | ="p469444" | done | 1 | 9 | Bacteria | 2 | cl21497~cl41762~NF041559 | PAAR_like~MIX~BTH_I2691_fam |
| 1 | ="p469444" | done | 1 | 7 | Pseudoalteromonas | 53246 | cl41762~cl47134~NF041559 | MIX~SMC_N~BTH_I2691_fam |
| 1 | ="p469444" | done | 1 | 7 | cellular organisms | 131567 | NF041559~pfam20455 | BTH_I2691_fam~DUF6708 |
| 1 | ="p469444" | done | 1 | 6 | Gilliamella | 1193503 | NF041559~COG3883 | BTH_I2691_fam~CwlO1 |
| 1 | ="p469444" | done | 1 | 6 | Gilliamella | 1193503 | NF041559~COG4372 | BTH_I2691_fam~COG4372 |
| 1 | ="p469444" | done | 1 | 6 | Gammaproteobacteria | 1236 | cl41762~TIGR00618~COG1196~NF041559 | MIX~sbcc~Smc~BTH_I2691_fam |
| 1 | ="p469444" | done | 1 | 6 | Bacteroides salyersiae | 291644 | COG3941~PTZ00121~NF041559 | HI1514~PTZ00121~BTH_I2691_fam |
| 1 | ="p469444" | done | 1 | 6 | Lachnospiraceae bacterium TM07-2AC | 2302966 | pfam20155~NF041559 | TMP_3~BTH_I2691_fam |
| 1 | ="p469444" | done | 1 | 5 | Vibrio | 662 | cl21497~COG4932~pfam18734~cl41762~NF041559 | PAAR_like~ClfA~HEPN_AbiU2~MIX~BTH_I2691_fam |
| 1 | ="p469444" | done | 1 | 5 | Bacteria | 2 | NF041559~PRK03918 | BTH_I2691_fam~PRK03918 |

| | | | | | | | | |
|---|---|---|---|---|---|---|---|---|
| 1 | ="p469444" | done | 1 | 5 | Vibrionaceae | 641 | cl21497~COG4932~NF041559 | PAAR_like~ClfA~BTH_I2691_fam |
| 1 | ="p469444" | done | 1 | 5 | Vibrionaceae | 641 | cl21497~COG4932~cl41762~NF041559 | PAAR_like~ClfA~MIX~BTH_I2691_fam |
| 1 | ="p469444" | done | 1 | 5 | Moritella | 58050 | COG1652~cl41762~NF041559 | XkdP~MIX~BTH_I2691_fam |
| 1 | ="p469444" | done | 1 | 4 | Bacteria | 2 | COG1388~NF041559 | LysM~BTH_I2691_fam |
| 1 | ="p469444" | done | 1 | 4 | Aeromonas | 642 | COG1388~NF041559~cl41762 | LysM~BTH_I2691_fam~MIX |
| 1 | ="p469444" | done | 1 | 4 | Gilliamella | 1193503 | NF041559~COG3883~cl15828 | BTH_I2691_fam~CwlO1~DUF308 |
| 1 | ="p469444" | done | 1 | 4 | Photobacterium | 657 | cl41762~cl21525~cl41762~NF041559 | MIX~LysM~MIX~BTH_I2691_fam |
| 1 | ="p469444" | done | 1 | 4 | Achromobacter xylosoxidans | 85698 | NF041559~COG3321 | BTH_I2691_fam~PksD |
| 1 | ="p469444" | done | 1 | 3 | Pseudomonadota | 1224 | COG3501~NF041559 | VgrG~BTH_I2691_fam |
| 1 | ="p469444" | done | 1 | 3 | Oceanospirillales | 135619 | cl21497~cl41762~cl21525~NF041559 | PAAR_like~MIX~LysM~BTH_I2691_fam |
| 1 | ="p469444" | done | 1 | 3 | Gammaproteobacteria | 1236 | COG3321~cl41762~NF041559 | PksD~MIX~BTH_I2691_fam |
| 1 | ="p469444" | done | 1 | 3 | unclassified Pseudomonas | 196821 | cl41758~NF041559 | peptidase_C58-like~BTH_I2691_fam |
| 1 | ="p469444" | done | 1 | 3 | unclassified Gilliamella | 2685620 | NF041559~pfam14282 | BTH_I2691_fam~FlxA |
| 1 | ="p469444" | done | 1 | 3 | Pseudoalteromonas | 53246 | cl41762~TIGR04523~NF041559 | MIX~Mplasa_alph_rch~BTH_I2691_fam |
| 1 | ="p469444" | done | 1 | 2 | Aeromonas | 642 | cl46249~NF041559~cl41762 | LysM~BTH_I2691_fam~MIX |
| 1 | ="p469444" | done | 1 | 2 | Streptococcus gordonii | 1302 | pfam20155~NF041559~COG3468 | TMP_3~BTH_I2691_fam~AidA |
| 1 | ="p469444" | done | 1 | 2 | Acinetobacter | 469 | cl00470~NF041559 | AKR_SF~BTH_I2691_fam |
| 1 | ="p469444" | done | 1 | 2 | Pseudomonadota | 1224 | cl21525~NF041559 | LysM~BTH_I2691_fam |
| 1 | ="p469444" | done | 1 | 2 | Pseudomonadota | 1224 | pfam13503~NF041559 | DUF4123~BTH_I2691_fam |
| 1 | ="p469444" | done | 1 | 2 | Gilliamella apicola | 1196095 | NF041559~PRK03918~PLN03229 | BTH_I2691_fam~PRK03918~PLN03229 |
| 1 | ="p469444" | done | 1 | 2 | Pseudoalteromonas luteoviolacea | 43657 | cl41762~cl47134~pfam11932~NF041559 | MIX~SMC_N~DUF3450~BTH_I2691_fam |
| 1 | ="p469444" | done | 1 | 2 | cellular organisms | 131567 | NF041559~COG2931 | BTH_I2691_fam~COG2931 |
| 1 | ="p469444" | done | 1 | 2 | Vibrio vulnificus | 672 | cl21497~pfam09595~cl41762~NF041559 | PAAR_like~Metaviral_G~MIX~BTH_I2691_fam |
| 1 | ="p469444" | done | 1 | 2 | Alcanivorax | 59753 | pfam10106~NF041559 | DUF2345~BTH_I2691_fam |
| 1 | ="p469444" | done | 1 | 2 | Bacillota | 1239 | pfam20155~NF041559~COG5412 | TMP_3~BTH_I2691_fam~COG5412 |
| 1 | ="p469444" | done | 1 | 2 | Pseudomonas citronellolis | 53408 | cl41762~pfam05335~NF041559 | MIX~DUF745~BTH_I2691_fam |

| | | | | | | | | |
|---|---|---|---|---|---|---|---|---|
| 1 | ="p469444" | done | 1 | 2 | unclassified Pseudoalteromonas | 194690 | cl41762~cl47134~pfam09727~NF041559 | MIX~SMC_N~CortBP2~BTH_I2691_fam |
| 1 | ="p469444" | done | 1 | 2 | unclassified Pseudomonas | 196821 | cl41762~pfam13002~NF041559 | MIX~LDB19~BTH_I2691_fam |
| 1 | ="p469444" | done | 1 | 2 | Gilliamella apicola | 1196095 | NF041559~cl47134~cl15828 | BTH_I2691_fam~SMC_N~DUF308 |
| 1 | ="p469444" | done | 1 | 2 | Gammaproteobacteria | 1236 | cl41762~TIGR00618~NF041559 | MIX~sbcc~BTH_I2691_fam |
| 1 | ="p469444" | done | 1 | 2 | unclassified Pseudoalteromonas | 194690 | cl41762~TIGR00606~TIGR04523~NF041559 | MIX~rad50~Mplasa_alph_rch~BTH_I2691_fam |
| 1 | ="p469444" | done | 1 | 1 | Thunnus | 8234 | NF041559~cl21478~cl02614 | BTH_I2691_fam~ATP-synt_Fo_b~SPRY |
| 1 | ="p469444" | done | 1 | 1 | Pseudomonas | 286 | cl41762~TIGR04320~NF041559 | MIX~Surf_Exclu_PgrA~BTH_I2691_fam |
| 1 | ="p469444" | done | 1 | 1 | Pseudomonas syringae group | 136849 | cl41762~cl09884~NF041559 | MIX~DUF2400~BTH_I2691_fam |
| 1 | ="p469444" | done | 1 | 1 | Butyrivibrio proteoclasticus | 43305 | COG0318~NF041559 | MenE/FadK~BTH_I2691_fam |
| 1 | ="p469444" | done | 1 | 1 | Pseudomonadota | 1224 | COG1652~cl46249~NF041559 | XkdP~LysM~BTH_I2691_fam |
| 1 | ="p469444" | done | 1 | 1 | Pseudoalteromonas | 53246 | cl41762~COG1579~PRK03918~NF041559 | MIX~DR0291~PRK03918~BTH_I2691_fam |
| 1 | ="p469444" | done | 1 | 1 | Pseudomonas chengduensis | 489632 | cl41762~COG0840~NF041559 | MIX~Tar~BTH_I2691_fam |
| 1 | ="p469444" | done | 1 | 1 | Gilliamella | 1193503 | NF041559~COG3883~PLN03229 | BTH_I2691_fam~CwlO1~PLN03229 |
| 1 | ="p469444" | done | 1 | 1 | unclassified Frigoribacterium | 2627005 | NF041559~COG0842 | BTH_I2691_fam~YadH |

### Dataset 2

| Accession | Description | Score | Coverage | # Proteins | # Unique Peptides | # Peptides | # PSMs | # AAs | MW [kDa] | calc. pI |
|---|---|---|---|---|---|---|---|---|---|---|
| **AAN68220.1** | **conserved membrane protein of unknown function** | 26570,76 | 73,29 | 1 | 72 | 72 | 647 | 996 | 109,7 | 7,46 |

| 10 | Sequence | # PSMs | # Proteins | # Protein Groups | Protein Group Accessions | Modifications | ΔCn | IonScore | Exp Value | Charge | MH+ [Da] | ΔM [ppm] | RT [min] | # Missed Cleavages |
|---|---|---|---|---|---|---|---|---|---|---|---|---|---|---|
| High | FPAmADITDLTAQSVNTVVLKR | 14 | 1 | 1 | DALBETKE | M4(Oxidation) | 0,0000 | 182 | 1,17495E-17 | 2 | 2406,23280 | -15,39 | 35,18 | 1 |
| High | YADAGFNLEQGFPTLQHSAYTLR | 15 | 1 | 1 | DALBETKE | | 0,0000 | 164 | 3,05726E-16 | 3 | 2599,22134 | -13,98 | 35,48 | 0 |
| High | FPAmADITDLTAQSVNTVVLK | 49 | 1 | 1 | DALBETKE | M4(Oxidation) | 0,0000 | 157 | 2,69025E-15 | 2 | 2250,13356 | -15,63 | 36,48 | 0 |
| High | ADSIGTLFGASDNVIEGLAGR | 13 | 1 | 1 | DALBETKE | | 0,0000 | 149 | 1,2681E-14 | 2 | 2063,01602 | -11,86 | 41,70 | 0 |
| High | DKADSIGTLFGASDNVIEGLAGR | 27 | 1 | 1 | DALBETKE | | 0,0000 | 147 | 3,00696E-14 | 2 | 2306,13396 | -12,34 | 40,20 | 1 |
| High | GPALSPAESHFQALTHEDYSPNGAR | 19 | 1 | 1 | DALBETKE | | 0,0000 | 140 | 4,28246E-14 | 3 | 2652,20728 | -13,78 | 26,55 | 0 |
| High | RLEQWLDQHDSPLYTALAPFNPFK | 5 | 1 | 1 | DALBETKE | | 0,0000 | 140 | 1,20761E-13 | 3 | 2886,42577 | -10,98 | 41,72 | 1 |
| High | LVIHEYVGAGLYKELR | 3 | 1 | 1 | DALBETKE | | 0,0000 | 137 | 2,83223E-13 | 3 | 1860,00253 | -19,00 | 28,95 | 1 |
| High | RGPALSPAESHFQALTHEDYSPNGAR | 19 | 1 | 1 | DALBETKE | | 0,0000 | 134 | 2,67361E-13 | 4 | 2808,30821 | -13,08 | 24,02 | 1 |
| High | DLmPAEQLEHTKPAQPPEQER | 17 | 1 | 1 | DALBETKE | M3(Oxidation) | 0,0000 | 130 | 1,0026E-12 | 3 | 2460,15265 | -12,13 | 18,63 | 0 |
| High | QTTPLQINPDcHITQAGLLLHYK | 8 | 1 | 1 | DALBETKE | C11(Carbamidomethyl) | 0,0000 | 128 | 2,61528E-12 | 3 | 2661,35200 | -11,20 | 31,42 | 0 |
| High | ALLAPYSEQLGLGALTTHLDINNK | 9 | 1 | 1 | DALBE | | 0,00 | 126 | 4,01541 | 2 | 2552,33 | - | 37, | 0 |

| | | | | | | | | | | | | | | |
|---|---|---|---|---|---|---|---|---|---|---|---|---|---|---|
| | | | | | TKE | | 00 | | E-12 | | 894 | 12,9 4 | 93 | |
| High | ITEQAITENPFSQEVQR | 13 | 1 | 1 | DALBE TKE | | 0,00 00 | 124 | 4,13306 E-12 | 2 | 1989,95 486 | - 16,5 0 | 30, 75 | 0 |
| High | mPLTWSSDPVTETLPIGNLHAIAR | 15 | 1 | 1 | DALBE TKE | M1(Oxidation) | 0,00 00 | 119 | 2,23624 E-11 | 3 | 2635,31 659 | - 14,5 7 | 35, 98 | 0 |
| High | SATLTINYYANQGLDDDTPSTTTIHLER | 14 | 1 | 1 | DALBE TKE | | 0,00 00 | 116 | 1,67918 E-11 | 3 | 3110,45 080 | - 13,0 4 | 32, 08 | 0 |
| High | ASGYPAIImGFSSDIFK | 13 | 1 | 1 | DALBE TKE | M9(Oxidation) | 0,00 00 | 116 | 2,68652 E-11 | 2 | 1819,87 526 | - 10,0 7 | 41, 08 | 0 |
| High | WPTPDWATIIHTQVTK | 16 | 1 | 1 | DALBE TKE | | 0,00 00 | 112 | 1,08075 E-10 | 3 | 1893,95 587 | - 15,8 2 | 36, 32 | 0 |
| High | GEKLVIHEYVGAGLYK | 1 | 1 | 1 | DALBE TKE | | 0,00 00 | 109 | 3,41962 E-10 | 3 | 1775,94 139 | - 15,6 1 | 24, 72 | 1 |
| High | GLEISLALLIHPmGQPTPGTEDHDRR | 10 | 1 | 1 | DALBE TKE | M13(Oxidation ) | 0,00 00 | 107 | 5,31253 E-10 | 3 | 2869,42 189 | - 14,2 0 | 34, 73 | 1 |
| High | QLDmAELVSGNQAPSTQPHVLPVSALQ TWVEDFKPTER | 4 | 1 | 1 | DALBE TKE | M4(Oxidation) | 0,00 00 | 103 | 7,65896 E-10 | 4 | 4235,05 949 | - 11,4 4 | 40, 55 | 0 |
| High | VPVVVALADAEGmALDLSLSVSAYQHQ LR | 6 | 1 | 1 | DALBE TKE | M13(Oxidation ) | 0,00 00 | 103 | 1,21747 E-09 | 3 | 3068,57 002 | - 12,5 9 | 42, 38 | 0 |
| High | GLEISLALLIHPmGQPTPGTEDHDR | 6 | 1 | 1 | DALBE TKE | M13(Oxidation ) | 0,00 00 | 102 | 1,14424 E-09 | 3 | 2713,32 625 | - 13,0 0 | 36, 38 | 0 |
| High | RmPLTWSSDPVTETLPIGNLHAIAR | 16 | 1 | 1 | DALBE TKE | M2(Oxidation) | 0,00 00 | 100 | 1,69986 E-09 | 3 | 2791,41 742 | - 13,8 6 | 33, 52 | 1 |
| High | mTNASVSSATSGPAcSAR | 2 | 1 | 1 | DALBE TKE | M1(Oxidation); C15(Carbamido methyl) | 0,00 00 | 98 | 2,64955 E-10 | 2 | 1770,75 998 | - 8,26 | 10, 72 | 0 |
| High | TLESLYPADAPSPELAAFR | 10 | 1 | 1 | DALBE TKE | | 0,00 00 | 97 | 2,88059 E-09 | 3 | 2048,00 077 | - 16,0 3 | 36, 18 | 0 |
| High | SDAPEYTLGFASSVFGVIGAAAATLVSV R | 11 | 1 | 1 | DALBE TKE | | 0,00 00 | 89 | 2,10644 E-08 | 2 | 2856,44 408 | - 11,8 5 | 42, 03 | 0 |
| High | ENDGEIRDNIKLDFDK | 4 | 1 | 1 | DALBE TKE | | 0,00 00 | 86 | 1,81199 E-08 | 3 | 1920,90 079 | - 15,1 4 | 25, 50 | 2 |
| High | LVIHEYVGAGLYK | 37 | 1 | 1 | DALBE TKE | | 0,00 00 | 86 | 6,14113 E-08 | 3 | 1461,78 091 | - 19,9 5 | 32, 88 | 0 |
| High | EANAEKALGLLAAR | 2 | 1 | 1 | DALBE TKE | | 0,00 00 | 85 | 3,77691 E-08 | 3 | 1426,77 802 | - 16,3 5 | 29, 22 | 1 |
| High | TVSIELTTTAR | 43 | 1 | 1 | DALBE | | 0,00 | 85 | 3,93673 | 2 | 1191,64 | - | 30, | 0 |

| | | | | | | | | | | | | | | |
|---|---|---|---|---|---|---|---|---|---|---|---|---|---|---|
| | | | | | TKE | | 00 | | E-08 | | 006 | 15,04 | 12 | |
| High | TNASVSSATSGPAcSAR | 6 | 1 | 1 | DALBETKE | C14(Carbamidomethyl) | 0,0000 | 85 | 1,38436E-08 | 2 | 1623,70970 | -18,16 | 10,30 | 0 |
| High | ENDGEIRDNIK | 8 | 1 | 1 | DALBETKE | | 0,0000 | 83 | 8,41724E-08 | 2 | 1302,61290 | -11,97 | 11,22 | 1 |
| High | LEQWLDQHDSPLYTALAPFNPFKDK | 5 | 1 | 1 | DALBETKE | | 0,0000 | 71 | 1,13048E-06 | 3 | 2973,45142 | -9,02 | 41,40 | 1 |
| High | SGmAQVEEmKALR | 4 | 1 | 1 | DALBETKE | M3(Oxidation); M9(Oxidation) | 0,0000 | 70 | 8,13803E-07 | 3 | 1481,68831 | -13,81 | 10,13 | 1 |
| High | LHSPLELWASR | 22 | 1 | 1 | DALBETKE | | 0,0000 | 70 | 1,31027E-06 | 2 | 1308,68434 | -16,51 | 32,65 | 0 |
| High | YRITEQAITENPFSQEVQR | 3 | 1 | 1 | DALBETKE | | 0,0000 | 69 | 1,00573E-06 | 3 | 2309,11702 | -15,21 | 30,53 | 1 |
| High | RLEQWLDQHDSPLYTALAPFNPFKDK | 3 | 1 | 1 | DALBETKE | | 0,0000 | 69 | 2,75188E-06 | 4 | 3129,54773 | -10,11 | 39,80 | 2 |
| High | ENDGEIRDNIKLDFDKK | 1 | 1 | 1 | DALBETKE | | 0,0000 | 67 | 2,33015E-06 | 3 | 2048,99578 | -14,18 | 22,13 | 3 |
| High | DNIKLDFDKK | 5 | 1 | 1 | DALBETKE | | 0,0000 | 66 | 5,20303E-06 | 2 | 1235,64294 | -16,31 | 17,38 | 2 |
| High | LDAEQLSPQSR | 7 | 1 | 1 | DALBETKE | | 0,0000 | 66 | 2,29698E-06 | 2 | 1243,61076 | -13,67 | 17,02 | 0 |
| High | DNIKLDFDK | 3 | 1 | 1 | DALBETKE | | 0,0000 | 64 | 4,33507E-06 | 2 | 1107,55020 | -16,18 | 22,35 | 1 |
| High | ALRPGYVYVFmK | 17 | 1 | 1 | DALBETKE | M11(Oxidation) | 0,0000 | 62 | 8,00085E-06 | 2 | 1459,75710 | -13,41 | 29,63 | 0 |
| High | ALRPGYVYVFMK | 1 | 1 | 1 | DALBETKE | | 0,0000 | 61 | 1,96224E-05 | 2 | 1443,76302 | -12,98 | 31,20 | 0 |
| High | NLALQASNDHDAWLATAEPQHIDNPYSLAAALAcYDR | 1 | 1 | 1 | DALBETKE | C34(Carbamidomethyl) | 0,0000 | 61 | 3,42027E-06 | 4 | 4095,88221 | -10,58 | 38,78 | 0 |
| High | NLVAmLLNK | 7 | 1 | 1 | DALBETKE | M5(Oxidation) | 0,0000 | 59 | 1,75201E-05 | 2 | 1031,57234 | -18,90 | 33,37 | 0 |
| High | SGmAQVEEmK | 13 | 1 | 1 | DALBETKE | M3(Oxidation); M9(Oxidation) | 0,0000 | 58 | 1,43277E-06 | 2 | 1141,47252 | -12,21 | 5,08 | 0 |
| High | YQGLEDYHR | 19 | 1 | 1 | DALBETKE | | 0,0000 | 57 | 4,05669E-06 | 3 | 1180,51708 | -17,88 | 14,95 | 0 |
| High | NLVAMLLNK | 1 | 1 | 1 | DALBETKE | | 0,0000 | 56 | 3,44923E-05 | 2 | 1015,57918 | -17,4 | 37,08 | 0 |

| | | | | | | | | | | | | | | |
|---|---|---|---|---|---|---|---|---|---|---|---|---|---|---|
| | | | | | | | | | | | | 7 | | |
| High | ALGLLAAR | 5 | 1 | 1 | DALBETKE | | 0,0000 | 55 | 2,79257E-05 | 2 | 784,48906 | -19,04 | 24,97 | 0 |
| High | SWQIEFFSPK | 15 | 1 | 1 | DALBETKE | | 0,0000 | 54 | 2,112E-05 | 1 | 1268,60872 | -17,60 | 37,17 | 0 |
| High | ITGSAALRK | 3 | 1 | 1 | DALBETKE | | 0,0000 | 53 | 7,38606E-05 | 2 | 916,54496 | -13,67 | 6,53 | 1 |
| High | YLKSGmAQVEEmK | 2 | 1 | 1 | DALBETKE | M6(Oxidation); M12(Oxidation) | 0,0000 | 53 | 1,82933E-05 | 3 | 1545,70153 | -17,66 | 8,77 | 1 |
| High | YQGLEDYHRR | 5 | 1 | 1 | DALBETKE | | 0,0000 | 53 | 3,07035E-05 | 3 | 1336,61845 | -15,60 | 11,35 | 1 |
| High | RQVQLAAR | 6 | 1 | 1 | DALBETKE | | 0,0000 | 51 | 8,15128E-05 | 2 | 941,54842 | -16,51 | 6,70 | 1 |
| High | TADAPAcR | 3 | 1 | 1 | DALBETKE | C7(Carbamidomethyl) | 0,0000 | 50 | 2,29731E-05 | 2 | 861,37770 | -12,39 | 4,60 | 0 |
| High | WDASHGLR | 10 | 1 | 1 | DALBETKE | | 0,0000 | 48 | 7,47254E-05 | 2 | 941,44284 | -16,98 | 14,68 | 0 |
| High | TAYNSLQK | 11 | 1 | 1 | DALBETKE | | 0,0000 | 47 | 0,000172294 | 1 | 924,46771 | -11,72 | 10,17 | 0 |
| High | LDFDKKLPPFR | 2 | 1 | 1 | DALBETKE | | 0,0000 | 47 | 0,000506205 | 3 | 1375,75042 | -16,65 | 26,23 | 2 |
| High | LDVDKYR | 6 | 1 | 1 | DALBETKE | | 0,0000 | 45 | 0,000365623 | 2 | 908,46874 | -16,40 | 13,42 | 1 |
| High | YAIVPR | 1 | 1 | 1 | DALBETKE | | 0,0000 | 44 | 0,001718447 | 2 | 718,41054 | -19,65 | 16,53 | 0 |
| High | ENDGEIR | 5 | 1 | 1 | DALBETKE | | 0,0000 | 44 | 3,59895E-05 | 2 | 832,36816 | -13,72 | 5,43 | 0 |
| High | ALLAPYSEQLGLGALTTHLDINNKWPTPDWATIIHTQVTK | 1 | 1 | 1 | DALBETKE | | 0,0000 | 41 | 0,000653761 | 5 | 4427,28919 | -11,47 | 41,55 | 1 |
| High | DLmPAEQLEHTKPAQPPEQERVPAcYR | 1 | 1 | 1 | DALBETKE | M3(Oxidation); C25(Carbamidomethyl) | 0,0000 | 41 | 0,000465753 | 4 | 3206,49633 | -12,34 | 21,18 | 1 |
| High | VPILPVR | 8 | 1 | 1 | DALBETKE | | 0,0000 | 40 | 0,000107249 | 2 | 793,51498 | -18,27 | 24,03 | 0 |
| High | LEQWLDQHDSPLYTALAPFNPFK | 1 | 1 | 1 | DALBETKE | | 0,0000 | 40 | 0,001349489 | 3 | 2730,31237 | -16,10 | 43,48 | 0 |
| High | NLALQASNDHDAWLATAEPQHIDNPYSLAAALAcYDRDER | 2 | 1 | 1 | DALBETKE | C34(Carbamidomethyl) | 0,0000 | 39 | 0,001967178 | 5 | 4496,04129 | -12,2 | 37,15 | 1 |

| | | | | | | | | | | | | | | |
|---|---|---|---|---|---|---|---|---|---|---|---|---|---|---|
| | | | | | | | | | | | | 1 | | |
| High | VPAcYR | 6 | 1 | 1 | DALBE TKE | C4(Carbamido methyl) | 0,00 00 | 39 | 0,00067 8586 | 2 | 765,361 42 | - 12,8 3 | 8,2 8 | 0 |
| High | KLPPFR | 1 | 1 | 1 | DALBE TKE | | 0,00 00 | 39 | 0,00324 7043 | 2 | 757,456 80 | - 20,0 1 | 13, 70 | 1 |
| High | HIHEEIKSWQIEFFSPK | 1 | 1 | 1 | DALBE TKE | | 0,00 00 | 37 | 0,00415 6813 | 4 | 2155,06 029 | - 17,1 2 | 31, 18 | 1 |
| High | LDFDKK | 1 | 1 | 1 | DALBE TKE | | 0,00 00 | 36 | 0,00298 4993 | 2 | 765,405 38 | - 11,4 8 | 10, 47 | 1 |
| High | FLAERDALER | 1 | 1 | 1 | DALBE TKE | | 0,00 00 | 35 | 0,00562 9453 | 3 | 1219,62 013 | - 18,7 8 | 17, 12 | 1 |
| High | ALRPGYVYVFmKGPR | 1 | 1 | 1 | DALBE TKE | M11(Oxidation ) | 0,00 00 | 35 | 0,00435 8302 | 4 | 1769,92 533 | - 15,0 9 | 23, 55 | 1 |
| High | QVQLAAR | 1 | 1 | 1 | DALBE TKE | | 0,00 00 | 32 | 0,00327 0904 | 2 | 785,449 58 | - 16,8 9 | 10, 03 | 0 |
| High | HYPWTQPK | 1 | 1 | 1 | DALBE TKE | | 0,00 00 | 30 | 0,00400 3764 | 2 | 1056,50 626 | - 18,8 3 | 14, 38 | 0 |

#### LEGENDS FOR SUPPLEMENTARY INFORMATION DATASETS

**Dataset S1. Conserved Domain Architecture of the BTH_I2691 protein family.** A dataset containing the list of proteins belonging to the BTH_I2691 protein family and identified using the Conserved Domain Architecture Retrieval Tool (CDART). Columns represent: A: index, B: query, C: status, D: score, E: number of sequences, F: taxonomy, G: taxid, H: the accession number of these proteins, and I: the name of the different domains identified in these proteins. The BTH_I2691 domain is found solo or in combination with many other domains.

**Dataset S2. Mass spectrometry identification of Tke5 purified band.** A dataset containing the list of proteins identified by mass spectrometry. Columns represent: A: accession number, B: description, C: score, D: coverage, E: proteins, F: unique peptides, G: peptides, H: PSMs, I: number of amino acids, J: molecular weight and K: isoelectric point.

#### SUPPLEMENTARY REFERENCES


Boyer, H.W. and Roulland-Dussoix, D. (1969) A complementation analysis of the restriction and modification of DNA in *Escherichia coli*. *J Mol Biol* **41**: 459–72.

Calles, B., Goñi-Moreno, Á., and Lorenzo, V. (2019) Digitalizing heterologous gene expression in Gram-negative bacteria with a portable ON/OFF module. *Mol Syst Biol* **15**: e8777.

Espinosa-Urgel, M., Salido, A., and Ramos, J.-L. (2000) Genetic Analysis of Functions Involved in Adhesion of *Pseudomonas putida* to Seeds. *J Bacteriol* **182**: 2363–2369.

Feil, H., Feil, W.S., Chain, P., Larimer, F., DiBartolo, G., Copeland, A., et al. (2005) Comparison of the complete genome sequences of *Pseudomonas syringae* pv. *syringae* B728a and pv. *tomato* DC3000. *Proc Natl Acad Sci U S A* **102**: 11064–11069.

González-Magaña, A., Tascón, I., Altuna-Alvarez, J., Queralt-Martín, M., Colautti, J., Velázquez, C., et al. (2023) Structural and functional insights into the delivery of a bacterial Rhs pore-forming toxin to the membrane. *Nat Commun* **14**: 7808.

Hanahan, D. (1985) Techniques for transformation of *E. coli*. In *DNA Cloning: A Practical Approach*. Glover, D.M. (ed). Virginia, p. 109.

Herrero, M., Lorenzo, V. de, and Timmis, K.N. (1990) Transposon vectors containing non-antibiotic resistance selection markers for cloning and stable chromosomal insertion of foreign genes in gram-negative bacteria. *J Bacteriol* **172**: 6557–6567.

Kessler, B., Lorenzo, V. de, and Timmis, K.N. (1992) A general system to integrate *lacZ* fusions into the chromosomes of gram-negative eubacteria: regulation of the *Pm* promoter of the TOL plasmid studied with all controlling elements in monocopy. *Mol Gen Genet* **233**: 293–301.

Kotoujansky, A., Diolez, A., Boccara, M., Bertheau, Y., Andro, T., and Coleno, A. (1985) Molecular cloning of *Erwinia chrysanthemi* pectinase and cellulase structural genes. *EMBO J* **4**: 781.

Martínez-García, E., Fraile, S., Algar, E., Aparicio, T., Velázquez, E., Calles, B., et al. (2023) SEVA 4.0: an update of the Standard European Vector Architecture database for advanced analysis and programming of bacterial phenotypes. *Nucleic Acids Res* **51**: D1558–D1567.

Nikel, P.I., Benedetti, I., Wirth, N.T., de Lorenzo, V., and Calles, B. (2022) Standardization of regulatory nodes for engineering heterologous gene expression: a feasibility study. *Microb Biotechnol* **15**: 2250–2265.

Pérez-Martínez, I., Rodriguez-Moreno, L., Matas, I., and Ramos, C. (2007) Strain selection and improvement of gene transfer for genetic manipulation of *Pseudomonas savastanoi* isolated from olive knots. *Res Microbiol* **158**: 60–69.

Silva-Rocha, R., Martínez-García, E., Calles, B., Chavarría, M., Arce-Rodríguez, A., Heras, A. de las, et al. (2013) The Standard European Vector Architecture (SEVA): a coherent platform for the analysis and deployment of complex prokaryotic phenotypes. *Nucleic Acids Res* **41**: D666–D675.

Studier, F.W. and Moffatt, B.A. (1986) Use of bacteriophage T7 RNA polymerase to direct selective high-level expression of cloned genes. *J Mol Biol* **189**: 113–130.
